# Massive reduction of RyR1 in muscle spindles of mice carrying recessive *Ryr1* mutations alters proprioception and causes scoliosis

**DOI:** 10.1101/2024.08.09.607317

**Authors:** Alexis Ruiz, Sofia Benucci, Hervé Meier, Georg Schultz, Katarzyna Buczak, Christoph Handschin, Rodrigo C. G. Pena, Susan Treves, Francesco Zorzato

## Abstract

Muscle spindles are stretch receptors lying deep within the muscle belly involved in detecting changes in muscle length and playing a fundamental role in motor control, posture and synchronized gait. They are made up of an external capsule surrounding 3-5 intrafusal muscle fibers and a nuclear bag complex. Dysfunction of muscle spindles leads to abnormal proprioceptor function, which has been linked to aberrant bone and cartilage development, scoliosis, kyphosis and joint contractures. *RYR1*, the gene encoding the calcium release channel of the sarcoplasmic reticulum, is the most common target of mutations linked to human congenital myopathies, a condition often accompanied by skeleton alterations and joint contractures. So far, the link between *RYR1* mutations, altered muscle spindles and skeletal defects has not been investigated. To this end, we investigated heterozygous mice carrying recessive *Ryr1* mutations isogenic to those present in a severely affected child. Here we show that: (i) the RyR1 protein localizes to the polar regions of intrafusal fibers and exhibits a doubled row distribution pattern, typical for junctional sarcoplasmic reticulum proteins; (ii) muscle spindles of compound heterozygous mice show structural defects; (iii) RyR1 content in intrafusal muscle fibers from dHT mice is reduced by 54%. Such a massive reduction of mutant RyR1 in intrafusal muscle fibers leads to altered expression of intrafusal fiber proteins, severe scoliosis, alteration of gait and inter limb coordination. These results support the hypothesis that *RYR1* mutations not only affect the function of extrafusal muscles, but might also affect that of intrafusal muscles. The latter may be one of the underlying causes of skeletal abnormalities seen in patients affected by recessive *RYR1* mutations.

## Introduction

Dysregulation of calcium homeostasis in skeletal muscles play an important role in the pathogenesis of a number of neuromuscular disorders (1, 2). Disruption of the calcium signals can be brought about by mutations in a number of genes encoding key proteins involved in excitation-contraction coupling (ECC) including the ryanodine receptor type 1 (RyR1), voltage-dependent calcium channels (dihydropyridine receptor, DHPR), STAC3, Stim1, Orai1 and calsequestrin 1 (CASQ1) (1–9). Depolarization of the sarcolemma is sensed by the DHPR which delivers the signal to release the Ca^2+^ stored in the sarcoplasmic reticulum (SR) terminal cisternae by opening the RyR1 calcium release channel, leading to muscle contraction by the process known as ECC. The re-uptake of cytosolic calcium by the sarcoplasmic reticulum CaATPase induces muscle relaxation (10–13). Mutations in *RYR1*, the gene encoding RyR1, underlie several neuromuscular disorders, including malignant hyperthermia (MH; MIM #145600), central core disease (CCD; MIM #11700), specific forms of multi-minicore disease and centronuclear myopathy (MmD, CNM; MIM # 255320) (1–4).

A great deal of experimental data has shown that *RYR1* mutations result in different types of channel defects ranging from alterations of regulatory mechanisms of channel activity to deficits of RyR1 expression (2, 14–16). The latter is characteristic to recessive mutations linked to congenital myopathies such as MmD and CNM. The clinical presentation of patients harboring recessive *RYR1* mutations usually occurs in infancy or early childhood and is characterized by motor developmental delays, hypotonia, proximal muscle weakness and muscle stiffness. Additional symptoms and phenotypes include involvement of extraocular muscles leading to ophthalmoplegia and/or ptosis and involvement of respiratory muscles, often requiring assisted ventilation (2–4,17–19). Of importance, many patients present a number of skeletal abnormalities at birth, including scoliosis and congenital dislocation of the hip, kyphosis, clubfoot (talipes equinovarus), flattening of the arch of the foot (flatfoot or pes planus), or an abnormally high arch of the foot (pes cavus) (4, 19). These observations are particularly intriguing and point to a potential role of the RyR1 in the correct development and maintenance of the musculoskeletal system, a hypothesis sustained by the observation that embryos of mice and birds depleted of RyR1, or carrying homozygous mutations in the *Ryr1* have skeletal abnormalities including kyphosis, excavated sternum, abnormal head positioning, spreading of the toes and crossing of the legs at the tibiotarsal joints (20–23).

Fetal skeletal muscle movements appear to be relevant for the appropriate development of the skeletal system and bone morphogenesis as demonstrated by the following observations: (i) fetal akinesia or insufficient fetal movements results in abnormal joint development and joint contractures (24, 25); (ii) the presence of *RYR1* mutations is also linked to fetal akinesia syndrome (26, 27); (iii) muscular dysgenesis mice carrying mutations in the α1s of the DHPR and lacking ECC, exhibit arrest of chondrocyte proliferation in the developing bone eminences, resulting in their loss (28).

Nevertheless, studies on the factors involved in appropriate musculoskeletal development are in their infancy. One recently identified player is the mechanosensitive ion channel Piezo2. In humans, mutations in *PIEZO2* lead to scoliosis, hip dislocation, joint contracture, arthrogryposis, respiratory distress and muscle weakness (29–33). Piezo2 is expressed in muscle spindles and Golgi tendon organs, two proprioceptive mechanosensory organs (34). Muscle spindles do not significantly contribute to the generation of muscle force but are stretch detectors. When a muscle is stretched, the change in length is transmitted to the muscle spindles which lie deep within the belly of the muscle; muscle spindles then convey the information concerning muscle stretch as well as the speed of stretching via sensory neurons to the central nervous system, which responds by adjusting the response of agonistic and antagonistic muscle groups (35). Each muscle spindle contains approximately 3-5 intrafusal muscle fibers (in mouse) made up of nuclear bag and nuclear chain fibers; the polar region of these specialized muscle fibers also contains contractile filaments as well as sarcotubular membranes including longitudinal tubules and triads similar to those present in extrafusal fibers (36, 37). Triads are endowed with calcium release units (CRU), and are formed by two SR terminal cisternae closely opposed to a central tubule oriented transversally (TT) with respect to the longitudinal axis of the myofibrils (12). In muscle spindles, activation of the CRU in response to gamma motor neuron stimulation, would cause contraction of the polar region of intrafusal fibers, an event which would offset the slackening of intrafusal fibers during the shortening of extrafusal fibers, thus providing a continuous adjustment of muscle spindle afferent firing during normal locomotor activity. On the basis of these findings, we hypothesize that the presence of mutated RyR1 in the CRU may affect the intrafusal fibers of muscle spindles by interfering with the adjustment of gamma motor neuron mediated intrafusal fiber contraction. This could result in proprioceptive signaling abnormalities in paravertebral muscles which in turn may induce musculoskeletal deformities, features that have been observed in many patients with *RYR1* mutations (3, 4, 19).

Here we investigated muscle spindle structure and gait properties in transgenic mice carrying the compound heterozygous RyR1 mutations RyR1p.Q1970fsX16+p.A4329D (referable as dHT) a model for recessive *RYR1* myopathies (38). We also investigated muscle spindle structure and gait properties of two additional mouse models carrying mutations in the *Ryr1*. In particular, we examined Ex36 mutant mice, carrying the heterozygous RyR1p.Q1970fsX16 mutation, leading to the expression of a single WT allele (39), and of Ho mice, carrying the homozygous RyR p.F4976L mutation (40). These two mouse models exhibit a milder skeletal muscle phenotype compared to the dHT model, as well as a smaller reduction of RyR1 expression and content (39, 40). Our results show that dHT mice, but not Ex36 and Ho mice, display changes in the histological appearance of muscle spindles and skeleton including: (i) alterations of the structure of the muscle spindles with an enlarged gap between the muscle spindle and extrafusal fibers, (ii) alterations of gait properties, (iii) skeleton abnormalities, in particular the presence of scoliosis as indicated by a 20% increase of the Cobb angle. High resolution proteomic analysis of muscle spindle samples collected by laser capture microdissection from WT and dHT mice confirm the presence of the RyR1 protein in intrafusal fibers and its strong reduction in content in muscle spindles from dHT mice compared to WT littermates.

In conclusion, our study shows that the presence of recessive RyR1 mutations might negatively influence not only the function of extrafusal muscles, but also that of muscle spindle intrafusal muscle fibers.

## Results

### RyR1 is expressed in intrafusal fibers

To demonstrate the presence of RyR1 in muscle spindles, we collected samples of intrafusal muscle fibers and extrafusal muscle fibers from WT mice by laser capture microdissection (LCM) and performed quantitative proteomic analysis of the LCM samples (Supplementary Fig.1). As experimental negative controls we also collected liver and kidney samples from WT mice and analyzed them using mass spectrometry. Figure 1A shows the PCA analysis of the proteomic data supporting the appropriateness of our experimental approach. In particular, the proteome of non-muscle tissues such as liver (Fig. 1A light brown samples) and kidney (Fig. 1A red samples) is very different from that of skeletal muscles (Fig. 1A green and violet samples) and the expression level of RyR1, if any, in liver and kidney is close to zero (Fig. 1B), confirming the specificity of the quantitative mass spectrometry analysis. Importantly, extrafusal muscle fibers (Fig.1A green samples) can be differentiated from intrafusal muscle fibers (Fig. 1A violet samples) by PCA analysis. Of note, extrafusal fibers contain 2.05-fold (p= 0.0028) more RyR1 protein compared to intrafusal fibers (Fig. 1B). On the basis of our previous quantitative mass spectrometry analysis of extrafusal fibers using spiked-in labelled peptides from RyR1 (41), the calculated average of RyR1 protein content in intrafusal fibres of WT mice is 0.628 µmol/kg of wet weight and the calculated RyR1 tetrameric complex is 0.152 µmol/kg wet weight.

**Figure 1:**
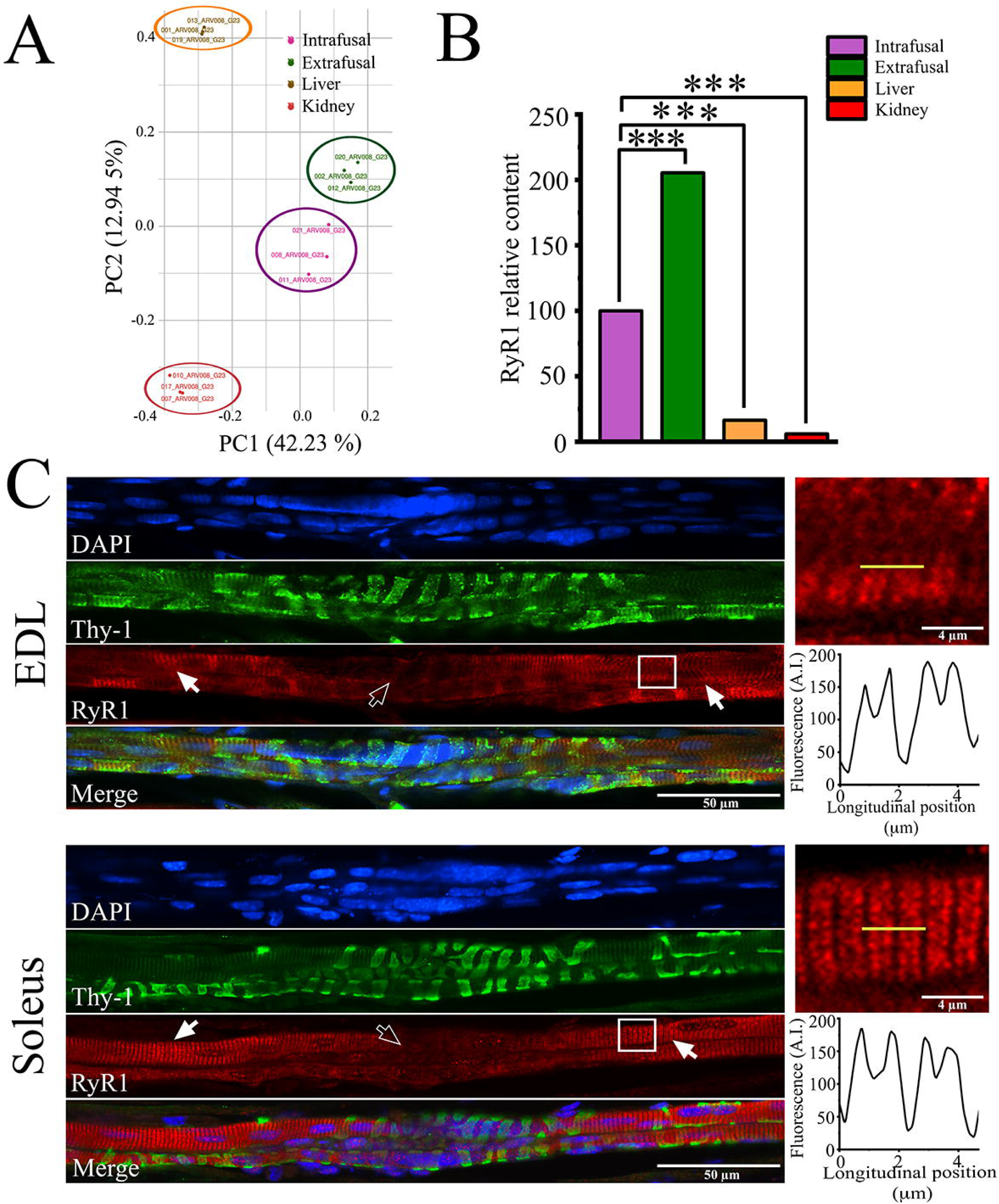
**Intrafusal muscle fibers express RyR1 protein and have a protein composition that is distinct from that of extrafusal muscle fibers**. **A.** PCA analysis of the proteome of intrafusal muscles (pink), extrafusal muscles (green), liver (brown) and kidney (red) from WT mice. Each symbol represents the analysis of proteins from one WT mouse (n = 3 mice). Intrafusal and extrafusal muscles were isolated form soleus muscles. **B.** Bar charts comparing relative RyR1 content in intrafusal fibers (set as 100%), extrafusal fibers, kidney and liver. RyR1 protein levels were obtained by mass spectrometry analysis on micro-dissected samples. Samples isolated from the intrafusal fibers, were selectively collected from the polar region of the fiber. Acquired reporter ion intensities in the experiments were employed for automated quantification and statistically analyzed using a modified version of our in-house developed SafeQuant R script (v2.3), the calculated p-values were corrected for multiple testing using the Benjamini−Hochberg method. **C.** Subcellular localization of RyR1 protein in muscle spindle fibers from 12 weeks WT mice. Confocal images were acquired along the longitudinal axis of muscle spindle fibers from EDL and soleus muscle. Images show the general structure of the muscle spindle, containing a polar region (white arrows) where RyR1 positive immunofluorescence is highest, and the central region (empty arrows) with a high content of nuclei (nuclear bag). Fibers were fixed and processed as described in the Methods section, and stained with DAPI (blue), Thy-1 (green) and RyR1 (red). Images were acquired using a Nikon AxR confocal microscope equipped with a 40x Plan Apo VC Nikon objective (NA= 0.95). Scale bars = 50 µm. The two insets show a higher magnification of the boxed region in EDL and soleus RyR1 staining, where the double labelling pattern characteristic of RyR1 distribution is clearly visible. White magnification bar = 4 µm. The yellow bar shows the longitudinal position of the fluorescently labelled RyR1. The lower panels of the two insets show the longitudinal spatial fluorescence profiles over the transverse direction within the line.

Having unequivocally shown that intrafusal fibers express an adequate amount of RyR1, we next investigated its subcellular localization by confocal immunohistochemistry. Longitudinal section of *extensor digitorum longus* (EDL) and Soleus muscles from Thy-1- EYFP transgenic mice were stained with anti-skeletal muscle RyR1 Ab and DAPI. Thy-1 marks all sensory neuron projections including group Ia and group II sensory afferent endings in the equatorial region of intrafusal muscle fibers (33). This area of intrafusal nuclear bag fibers is enriched in nuclei as indicated by DAPI staining. (Fig. 1C, left panels). The equatorial region extends into the polar region. The RyR1 Ab preferentially stains the polar regions of the intrafusal fibers (white arrows) but not the central nuclear bag region (empty arrows). The density profile of the RyR1 Ab staining shows the doubled row pattern typically observed in proteins located in the junctional SR of extrafusal fibers (42) (Fig.1C, right panels). No apparent difference in DAPI or RyR1 staining was observed between the muscle spindles of EDL and soleus muscles (Fig. 1C)

### Proteomic analysis reveals significant differences in the protein composition of intrafusal versus extrafusal muscles

The protein content of intrafusal and extrafusal fibers was analyzed under stringent conditions by removing proteins having ≤2 peptides, exhibiting a fold change ≥0.20 (- 0.321≤Log2FC≤0.263) and a q value ≤0.05. By filtering the mass spectrometry results using these stringent parameters, our results show that intrafusal muscle fibers have a significantly different protein composition compared to extrafusal fibers showing 542 down- regulated and 1591 up-regulated proteins (Fig. 2A). GO pathway analysis of the up- regulated and down-regulated proteins in intrafusal versus extrafusal muscle fibers identified proteins preferentially clustering to biological processes, molecular function and local network (Fig.2B and 2C). Within the biological process category, the pathways showing the largest enrichment score of up-regulated genes belong to the filopodium assembly, astrocyte development and xenobiotic transport groups. Within the molecular function category, the pathways showing the largest enrichment score of up-regulated genes belong to the xenobiotic membrane transport activity, fibronectin binding and extracellular matrix binding groups. Within the local network cluster category, the pathways showing the largest enrichment score of the up-regulated genes belong to the neuregulin binding, integrin alphav complex, aggrecan/versican and amino acid transport groups (Fig. 2B). As to the down-regulated pathways, within the biological process category, the pathways showing the largest down-regulated enrichment score belong to the electron transport coupled proton transport, fatty acid ß-oxidation and mitochondrial electron transport. Within the molecular function category, the pathways showing the largest down- regulated enrichment score belong to cytochrome c oxidase activity, oxidoreduction-driven transmembrane transport and ubiquinone binding. Within the local network cluster category, the pathways showing the largest down-regulated enrichment score belong to phosphorylase kinase complex, Hsp20/α crystallin family and FATZ binding and detection of muscle stretch (Fig. 2C). Table 1 shows the 50 top proteins showing the greatest fold change in intrafusal versus extrafusal muscles from WT mice. Cardiac troponin T (TnnT2), Somatomedin B and Thrombospondin Type 1 Domain Containing protein (Sbspon), dipeptidyl peptidase 4 (Dpp4), Monooxygenase DBH Like 1 (Moxd1), integrin ß4 subunit (Itgb4) and collagen type XXVIII Alpha 1 Chain (Col28a1) being the most enriched in intrafusal muscle fibers and Nicotinamide nucleotide transhydrogenase (Nnt), myosin-binding protein C2 family (Mybcp2), aquaporin 7 (Aqp7), Phosphorylase b kinase gamma catalytic chain (Phkg1) and Trio Rho Guanine Nucleotide exchange factor (Trio) being the least enriched in intrafusal muscles from WT mice. The Mass spectrometry data has been deposited in the ProteomeXchange Consortium via the ProteoSAFe repository (http://massive.ucsd.edu/ProteoSAFe) with the following access number PXD054222 (https://massive.ucsd.edu/ProteoSAFe/private-dataset.jsp?task=09ff33af463e430f8e40ce18bf5cf1d2).

**Figure 2:**
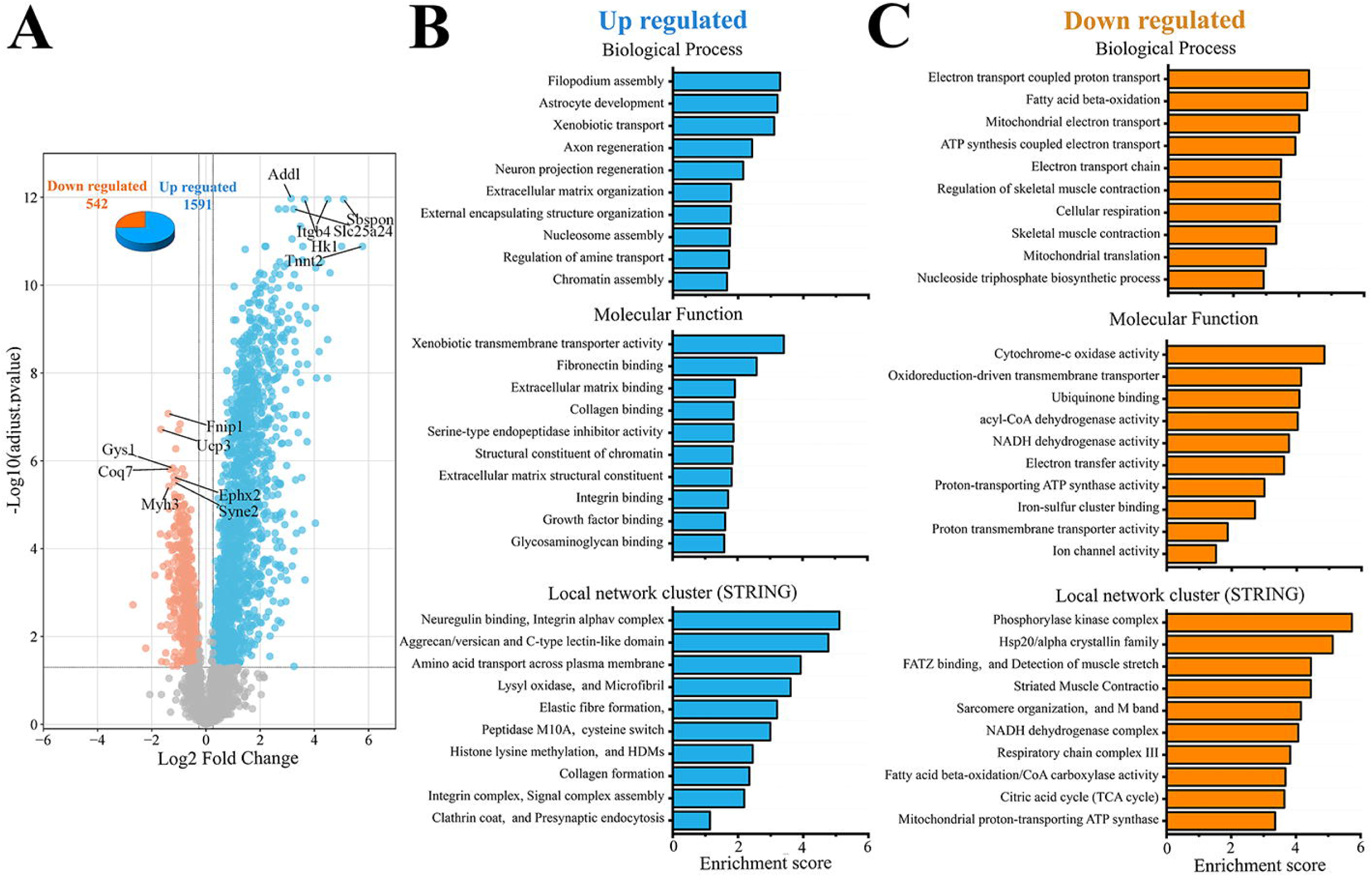
Comparison of the protein composition of intrafusal vs extrafusal muscle fibers from WT mice. A. Volcano plot analysis comparing protein content of intrafusal and extrafusal muscle fibers from soleus muscles of WT mice. The data was filtered so that only proteins quantified based on 2 ≥ peptides, exhibiting fold change ≥0.20 (- 0.321≤Log2FC≤0.263) and a q value ≤0.05 were considered as differentially expressed. More than 500 proteins were down-regulated in intrafusal vs extrafusal muscle fibers whereas more than 1500 were enriched in intrafusal versus extrafusal muscle fibers. **B.** Proteins showing a significant enrichment in intrafusal vs extrafusal muscles were analyzed by GO Pathway analysis. **C.** Proteins showing a significant depletion in intrafusal vs extrafusal muscles were analyzed by GO Pathway analysis. In both panels **B** and **C**, the enrichment scores were calculated based on Log_2_ FC for protein quantification based on 2 ≥ peptides. Data are plotted as Enrichment score vs significantly altered GO pathways. Muscles were collected from 5 WT and 5 dHT mice.

**Table 1:** Top 50 proteins showing the greatest fold change in content in intrafusal versus extrafusal muscle fibers in WT mice.

| Top 50 Up-regulated |  |  | Top 50 Down-regulated |  |  |
| --- | --- | --- | --- | --- | --- |
| Gene Name | Fold Change | q value | Gene Name | Fold Change | q value |
| Tnnt2 | 54.785 | 0.019921 | Nnt | 0.1547 | 1.11E-12 |
| Sbspon | 33.795 | 0.001899 | Mybpc2 | 0.2149 | 1.31E-11 |
| Dpp4 | 32.134 | 0.018494 | Aqp7 | 0.2719 | 5.25E-11 |
| Moxd1 | 23.913 | 0.000402 | Phkg1 | 0.2869 | 1.11E-12 |
| Itgb4 | 22.624 | 1.08E-05 | Trio | 0.3120 | 1.74E-09 |
| Col28a1 | 22.486 | 4.63E-05 | Abra | 0.3158 | 1.31E-08 |
| Sgce | 22.362 | 0.005811 | Ucp3 | 0.3169 | 2.97E-11 |
| Atp1b3 | 19.165 | 1.92E-07 | Tpm1 | 0.3225 | 3.26E-09 |
| Postn | 18.099 | 0.037256 | Tmod4 | 0.3355 | 1.30E-08 |
| Fbln2 | 16.913 | 0.000254 | Myl2 | 0.3582 | 4.01E-11 |
| Slc43a1 | 16.609 | 0.023672 | Mtx2 | 0.3614 | 3.32E-10 |
| Col18a1 | 16.458 | 0.038686 | Pak1 | 0.3706 | 2.63E-05 |
| Cend1 | 16.439 | 0.005393 | Cox7a1 | 0.3713 | 7.39E-10 |
| Akap12 | 13.495 | 5.18E-05 | Cox7b | 0.3727 | 6.31E-09 |
| Fbln1 | 13.388 | 0.000219 | Fnip1 | 0.3809 | 3.66E-07 |
| Olfml3 | 13.298 | 8.39E-08 | Fabp3 | 0.3823 | 9.15E-08 |
| Slc4a4 | 12.962 | 0.00043 | Fsd2 | 0.3863 | 3.06E-10 |
| Kif21a | 12.817 | 8.23E-05 | Fbp2 | 0.3875 | 0.00051582 |
| Thsd4 | 12.621 | 0.000338 | Myh3 | 0.3884 | 1.11E-12 |
| Hk1 | 12.454 | 3.82E-06 | Retreg1 | 0.3892 | 1.31E-11 |
| Itga6 | 11.919 | 1.24E-05 | Pxmp2 | 0.3916 | 2.69E-11 |
| Padi2 | 11.865 | 0.001573 | Apobec2 | 0.3952 | 3.78E-05 |
| Myl4 | 11.700 | 4.59E-05 | Coq7 | 0.3956 | 2.40E-06 |
| Hapln1 | 11.613 | 1.54E-06 | Ddo | 0.3975 | 1.06E-12 |
| Add3 | 11.265 | 0.000416 | Prr33 | 0.4062 | 1.64E-08 |
| Igfbp6 | 11.232 | 0.002048 | Hspb7 | 0.4067 | 4.58E-12 |
| Hspa12a | 11.194 | 0.003513 | Myoz2 | 0.4083 | 6.21E-10 |
| Lpar1 | 11.041 | 0.001682 | Hspb6 | 0.4102 | 1.16E-10 |
| Abcb1a | 10.720 | 0.007356 | Etfrf1 | 0.4103 | 6.31E-09 |
| Htra1 | 10.715 | 8.35E-05 | Ndufb8 | 0.4152 | 2.28E-07 |
| Fzd2 | 10.606 | 0.000301 | Myh8 | 0.4198 | 4.72E-10 |
| Atp1a3 | 10.479 | 0.000111 | Casq1 | 0.4270 | 4.45E-05 |
| Enpp1 | 10.098 | 0.046816 | Gys1 | 0.4274 | 8.99E-08 |
| Igsf8 | 10.082 | 1.46E-06 | Flnc | 0.4327 | 9.74E-09 |
| Xpnpep2 | 10.070 | 0.000205 | Phka1 | 0.4372 | 2.73E-06 |
| Rab34 | 9.911 | 0.000105 | Mylpf | 0.4387 | 3.32E-09 |
| Isyna1 | 9.818 | 0.005342 | Ephx2 | 0.4398 | 1.92E-09 |
| Col12a1 | 9.758 | 2.34E-06 | Des | 0.4403 | 5.73E-11 |
| Sfxn3 | 9.610 | 1.02E-05 | Mief2 | 0.4413 | 1.63E-05 |
| Atp6v1g2 | 9.576 | 0.002403 | Acsl1 | 0.4417 | 0.04790694 |
| S100a6 | 9.520 | 3.25E-05 | Ubr3 | 0.4424 | 1.85E-12 |
| Slc25a24 | 9.477 | 0.02924 | Scn4a | 0.4432 | 0.00011362 |
| Clec3a | 9.465 | 0.00072 | Perm1 | 0.4438 | 1.03E-09 |
| IGHG3 | 9.263 | 0.000854 | Acad11 | 0.4441 | 8.91E-08 |
| Igfbp5 | 9.227 | 0.00139 | Aqp1 | 0.4448 | 2.61E-11 |
| Sntb1 | 9.205 | 0.002906 | N4bp1 | 0.4454 | 0.00024887 |
| Smarcd3 | 9.110 | 0.002818 | Syne2 | 0.4463 | 8.87E-12 |
| Epb41l2 | 9.057 | 3.06E-06 | Acadvl | 0.4490 | 1.06E-12 |
| Add1 | 8.841 | 5.84E-06 | Higd1a | 0.4502 | 1.02E-10 |
| Itih5 | 8.781 | 0.001438 | Myot | 0.4503 | 9.05E-5 |

We also compared the data from the present investigation to recent transcriptional and proteomic data of muscle spindles versus extrafusal muscles (43, 44) from WT mice. Our results confirm that muscle spindles are enriched in a number of proteins including acetylcholinesterase (fold change= 1.676), collagen 3α1 and 6α1 (fold change= 2.06 and 2.299 respectively), α subunit of cardiac myosin heavy chain (fold change= 7.029), immunoglobulin-like and fibronectin type III domain containing 1 (fold change= 4.228), high mobility group protein B1 (fold change= 2.535), versican (fold change= 5.291), elastin (fold change= 3.610). fibulin-3 (fibrillin like protein, fold change 6.485), periostin (fold change= 18.099), Na+/K+ ATPase 3 (fold change= 10.479), heat shock 70 kD protein 12A (fold change= 11.194), vesicle amine transport-1-like (fold change= 3.119), myelin proteolipid protein 1 (fold change= 6.570), ß- tubulin 3 (fold change= 4.594), Peptidyl Arginine Deiminase 2 (fold change= 11.865) and myosin light chain 4 (fold change= 11.7) (Supplementary Table 1). Furthermore, Thy1 which we used as a marker to identify muscle spindles in immunohistochemical confocal experiments, was enriched 6.624-fold.

In summary, proteomic analysis unambiguously demonstrates that RyR1 is expressed in intrafusal muscle fibers, along with a large set of other proteins, leading us to next investigated whether RyR1 mutations affect muscle spindles in a murine model for a RyR1 linked recessive congenital myopathy.

### Expression of compound heterozygous mutant RyR1 in intrafusal fibers affects the gross histological appearance of muscle spindles

In order to investigate whether the presence of the RyR1p.Q1970fsX16+p.A4329D compound heterozygous mutations affect the gross morphological appearance of intrafusal muscle fibers, we performed serial sections of soleus muscles isolated from 12 weeks old WT and dHT mice. Figure 3A shows a schematic representation of the experimental approach and Fig. 3B shows representative H&E-stained sections of isolated muscles from WT and dHT mice. Once a muscle spindle was clearly identified in the section (Fig. 3A and B_a) the subsequent sections were made at a distance of 50 µm and then every 10 µm until the spindle was no longer visible (Fig.3A and B). The spindles of both WT and dHT mice (empty arrowhead) are embedded within, and surrounded by, extrafusal fibers (white arrowhead), however, it is clear that intrafusal muscle fibers of dHT mice are surrounded by a wider gap between the muscle spindle and the extrafusal fibers compared to those of WT mice. The maximal diameter (equatorial region) of the muscle spindle was identified by scrolling through the individual stack images (45) and was calculated as described in the Methods section. The mean (± SD) maximal diameter of muscle spindles of dHT mice was 71.2 ± 5.8 µm (n=15 spindles from n=5 mice) and larger than that of spindles of WT mice (60.6 ± 7.2 µm) (n=8 spindles from n= 3 mice) (p= 0.0197 Mann Whitney) (Fig.3C). However, the number of spindles per muscle and their location within the muscle belly, did not differ between WT and dHT mice. Furthermore, the mean (± S.D) number of muscle spindles in the belly of soleus muscles from WT and dHT mice was 3.5±0.4 (n=6) and 3.6±1.3 (n=11), respectively (p=0.120 Mann Whitney test). In muscle spindles from WT and dHT mice the number of nuclear bag and chain fibers was 1.90±0.29 and 1.83±0.38 (mean± S.D., n= 18, p=0.491 Mann Whitney) and 3.72±0.88 and 3.88±0.83 (mean± S.D., n=22, p= 0.676.

**Figure 3:**
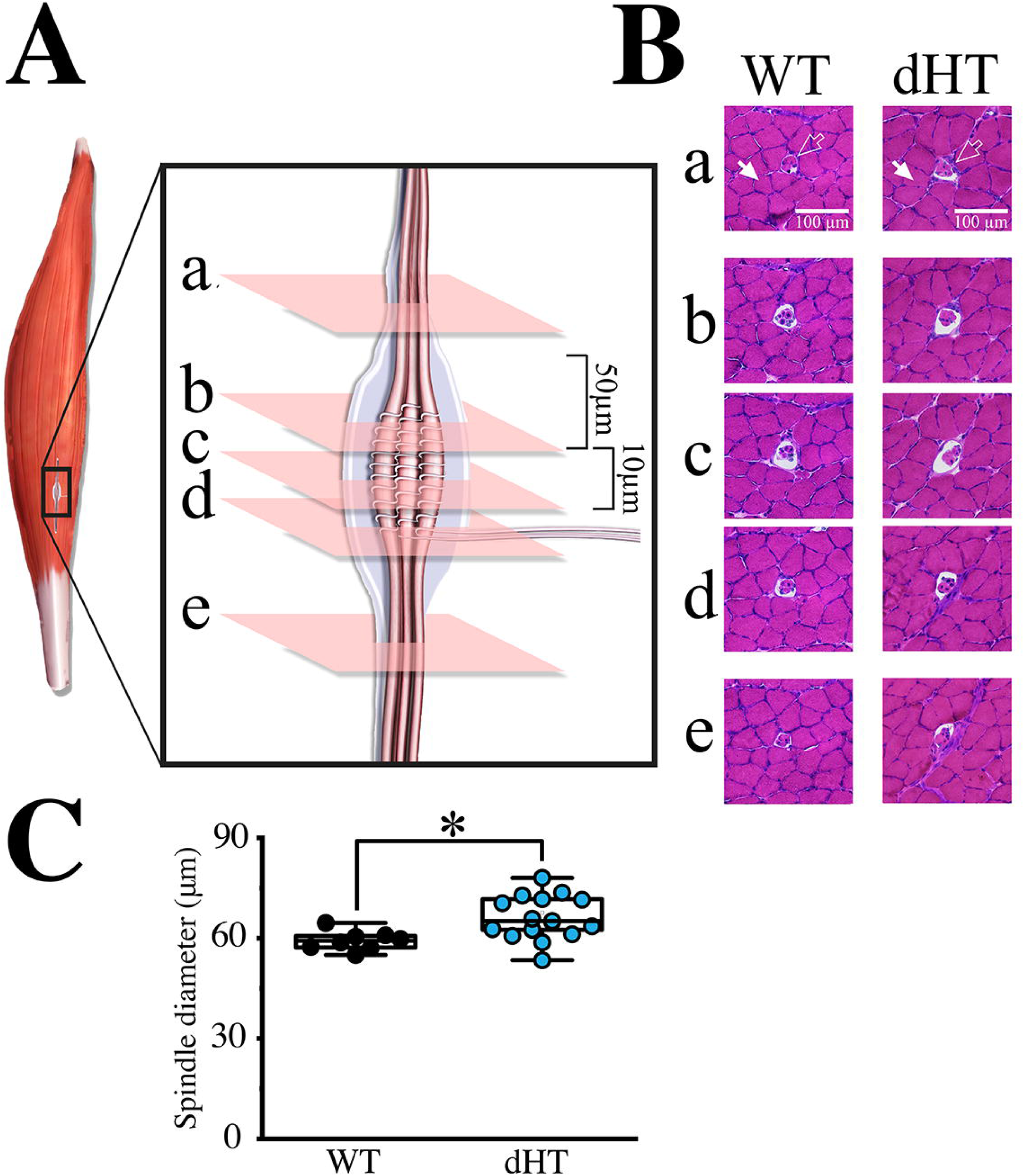
dHT mice exhibit gross alterations of muscle spindle morphology. A. Schematic representation of a skeletal muscle showing the location of a muscle spindle. Skeletal muscles contain several longitudinally oriented muscle spindles ubiquitously distributed in the interior of the muscle belly. They are composed of intrafusal fibers (longitudinally oriented pink fibers) surrounded by a capsule (represented as a light blue matrix). Transverse planes (“a”, “b”, “c”, “d” and “e”) were made through the muscle spindles at 5 different levels, “a” and “e” are located in the polar regions, “b”, “c” and “d” are located in the central region. **B.** H&E staining of soleus muscles sections containing intrafusal and extrafusal fibers from WT and dHT mice in. Ten µm transversal muscle sections were made until the polar region was identified. Extrafusal fiber are indicated by white arrows, intrafusal fibers are indicated by empty arrows. White bar in panels “a” = 100 µm. Images were acquired with an Eclipse Ti2 Nikon widefield microscope**. C.** Whisker plots of muscle spindle diameter. The diameter was calculated in the equatorial section of the muscle spindle represented as region “c” in panels A and B. The spindle diameter (distance of parallel tangents at opposing borders of the fiber was calculated using Fiji. Each symbol represents results obtained from a single spindle fiber (8 spindles from n= 3 WT mice; 15 spindles from dHT n=5 mice). *p < 0.05 (Mann-Whitney test).

In order to determine whether the altered histological appearance of the muscle spindles is a consequence of the RyR1p.Q1970fsX16+p.A4329D compound heterozygous mutations and/or occurs in the presence of other RyR1 mutations, we also examined H&E stained muscles from transgenic Exon 36 mice, which were used to generate the dHT mice. Mann Whitney), respectively. These carry a frameshift mutation leading to a down- stream premature stop codon causing nonsense mediated RNA decay of the mutant allele and resulting in ≈40% decrease of RyR1 content (39). Additionally, we included muscles from Ho mice, a transgenic mouse line carrying the homozygous RyR1 mutation p.F4976L identified in a severely affected child (40) (see Supplementary Figure 2 for a summary of the phenotypic characteristics of the transgenic mice used in the present investigation). As shown in Supplementary Fig.3, the gross morphological appearance and spindle diameters were not significantly different between Ho, Ex36 and WT mice.

In conclusion these results show that in mice carrying the compound heterozygous RyR1p.Q1970fsX16+p.A4329D mutations, but not those carrying the heterozygous Exon 36 mutation, nor the homozygous RyR1 p.F4976L mutation, the gross morphological appearance of muscle spindles is altered. We next performed behavioral tests aimed at verifying whether the morphological alterations of muscle spindles observed in the compound heterozygous RyR1p.Q1970fsX16+p.A4329D mice are paralleled by changes in proprioceptive properties such as gait and interlimb coordination.

### Gait and interlimb coordination properties are compromised in dHT mice

We investigated if the presence of the RyR1p.Q1970fsX16+p.A4329D compound heterozygous mutations affects balance or causes changes of the gait, by examining the capacity of age and sex-matched WT and dHT littermates to perform the Beam walk and the Catwalk, two behavioural tests that have been shown to be altered in a variety of transgenic mouse models including models with impaired muscle spindle function (46, 47). For the Beam walk test, mice were scored for their ability to walk across a 1 m long, 1 cm wide beam suspended on two poles 50 cm above a table top without losing their balance and for their capacity to precisely place their hindlimbs, as described in the Methods section and in Supplementary Figure 4A. All tests were scored by genotype blinded experimenters. Videos 1 and 2 and Figure 4A show that dHT mice (n=13) slip when walking across the beam, whereby they have a significantly higher Beam walk score than their WT (n=8) littermates (the Beam walk score was 0.086± 0.07 and 1.04±0.40 for WT and dHT, respectively p= 0.0014 Mann Whitney test). No significant differences were observed in the Beam walk score between WT and Ex 36 littermates (0.086± 0.07 and 0.071±0.220 for WT and Ex36, respectively p= 0.18 Mann Whitney test) and WT and Ho littermates (0.067± 0.133 and 0.083±0.108 for WT and Ho, respectively (p= 0.45 Mann Whitney test) (Supplementary Figure 4B). Since all values obtained on Ho mice were similar to those of WT littermates, we did not evaluate further Ho mice.

**Figure 4:**
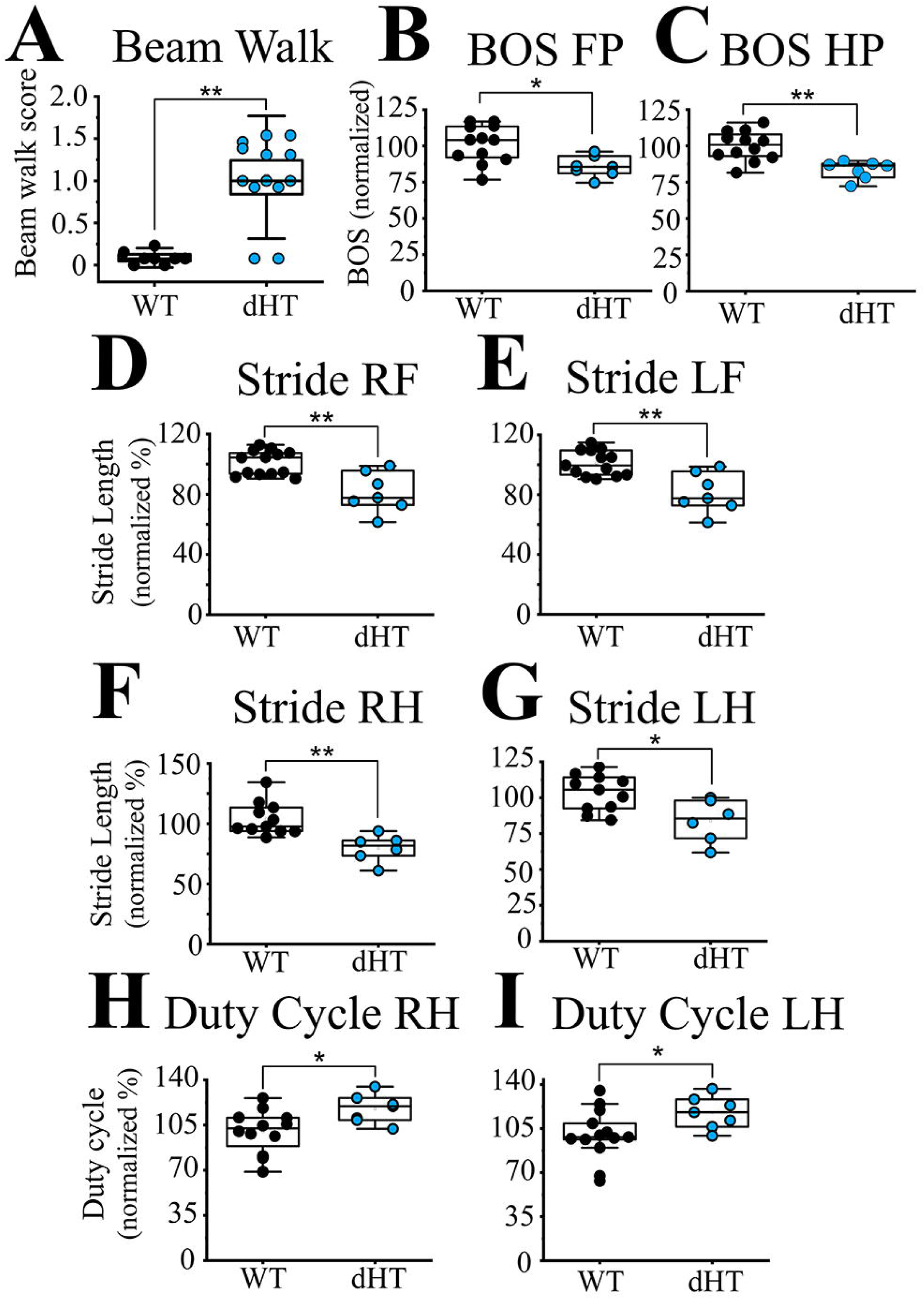
dHT mice show proprioceptor function alterations. A. The beam walk score test of WT mice is significantly lower than that of dHT littermates. Each symbol represents results obtained from a single mouse: WT (n= 8), dHT (n= 13) **p< 0.01 (Mann-Whitney test). **B-I** Catwalk gait analysis test performed using the CatWalk XT system. Twelve weeks old mice were acclimatized and studied on 3 consecutive days. Each day 3 successful runs were averaged per mouse. **B.** Base of support, Front Paws (BOS FP). **C.** Base of support, Hind Paws (BOS HP). **D.** Stride Length, Right Front Limb (Stride RF). **E.** Stride Length, Left Front limb (Stride LF). **F.** Stride Length, Right Hind limb (Stride RH). **G.** Stride Length, Left Hind limb (Stride LH). **H.** Duty Cycle, Right Hind (Duty Cycle RH). **I.** Duty Cycle, Left Hind (Duty Cycle LH). Each symbol represents results obtained from a single muscle. WT, n= 12 and dHT n= 7. *p < 0.05, **p< 0.01 (Mann-Whitney test). Results are normalized to values obtained in WT mice (=100%).

The Catwalk is an automated gait analysis system allowing the observer-independent and simultaneous quantification of multiple gait parameters (see Supplementary Fig. 5 for an explanation of the assessed parameters) as well as temporal aspects of interlimb co-ordination (46, 48). Due to the smaller size and weight of dHT mice, data was normalized to body weight to avoid experimental artefacts. Some CatWalk parameters did not differ between WT and dHT mice, including kinetic parameters and velocity, however, parameters reflecting interlimb coordination, were different in dHT compared to WT littermates (Fig. 4B-I and Supplementary Table 2). Within the interlimb coordination parameters dHT mice had a significantly lower Base of Support (BOS, average width of paws) of front and hind paws (Fig. 4B and 4C). Specifically, the BOS of the front paws (normalized, mean ± SD) of WT and dHT was 100.0±12.6 vs 81.2±7.2, respectively (p= 0.012, Mann Whitney test). The hind paws BOS (normalized, mean ± SD) of WT and dHT was 100.0±9.2 vs 83.4±6.2, respectively (p= 0.0027, Mann Whitney test). We also analyzed the Stride Length a parameter which detects gait alterations (49), by measuring the distance between successive placements of the same paw, measured during maximal contact. In the stride, both, front and hind paws of dHT mice showed a significant decrease compared to those of WT littermates, (Fig. 4D-G). The stride length, was 20 % lower in the four limbs of dHT mice, Right front (WT= 100.0±7.6 vs dHT 81.2±13.3; p= 0.0089 Mann Whitney test), Left front (WT= 100±7.7 vs dHT 81.1±13.2; p= 0.0071 Mann Whitney test), Right hind (WT= 100.0±11.9 vs dHT 79.6±11.5, p= 0.003, Mann Whitney test) and Left hind (WT= 100.0± 11.5 vs dHT 83.8±15.0; p= 0.023, Mann Whitney test). Finally, the Duty cycle defined as percentage of Stand (duration in seconds of contact of a paw with the glass plate) was also significantly higher in dHT mice compared to WT littermates. Specifically, the Right hind (normalized, mean ± SD) of WT and dHT was 100.0±16.6 vs 123.2 ±11.7, respectively (p= 0.0031, Mann Whitney test). While the Left hind (normalized, mean ± SD) of WT and dHT were 100.0 ±16.5 vs 117.6 ±12.9, respectively (p=0.039; Mann Whitney test). None of the examined parameters except for the Right hindlimb stand which was increased in Ex36 mice (134.9±29.3 p= 0.026), differed between WT and Ex36 and (Supplementary Figure 4 and Supplementary Table 2).

In conclusion, the CatWalk analysis results demonstrate that the scores of WT mice are significantly different from those of dHT littermates, supporting our hypothesis that the compound heterozygous RyR1p.Q1970fsX16+p.A4329D mutations not only affect extrafusal muscles, but also affect proprioceptive functions controlled by muscle spindles. In the next experiments, we investigated whether the compound heterozygous RyR1p.Q1970fsX16+p.A4329D mutations affect spine alignment, another indicator of proper proprioceptor function (50).

### dHT mice show skeleton abnormalities

The proprioceptive system is involved in maintaining spinal alignment and indeed, patients with recessive *RYR1* myopathies often present several skeleton abnormalities from birth as well as joint contractures. In addition, skeleton abnormalities occur in murine models lacking Egr3 or RunX3, key proteins which are needed for the proper function of muscle spindle and proprioceptive circuitry, respectively (50). On the basis of these observations, we next analyzed by high resolution microtomography whether dHT mice show skeleton abnormalities. Figure 5A shows a representative image of the whole-body view of the skeletons of WT and dHT 12 weeks old mice. The red line indicating the trajectory of the spinal cord confirm that dHT exhibit scoliosis and the white arrows mark changes of the spinal alignment. In order to quantify the degree of scoliosis, we measured Cobb angle values, which are the most widely used measurements to quantify the magnitude of scoliosis. Figure 5B shows Cobb angle values for the spine of WT and dHT mice and each symbol represents results obtained from a single mouse (n=6 WT and n=6 dHT). The spine of dHT mice have a calculated mean±SD Cobb angle of 19.3± 9.4 whereas the spines of WT littermates have a mean Cobb angle of 3.6±2.5 p=0.036, Mann-Whitney test). While examining the skeletons, we also observed marked changes in the spinal alignment leading to kyphosis (Fig. 5C) in dHT mice (see https://zenodo.org/records/12724636 for complete dataset of skeleton analysis). On the other hand, Ex36 mice showed no significant skeleton alterations; the mean± SD Cobb angle values for the spine were 2.3±0.9 (n=5 mice) versus 2.6±0.6 (n=5 mice), in WT and Ex36, respectively (Supplementary Fig. 6).

**Figure 5:**
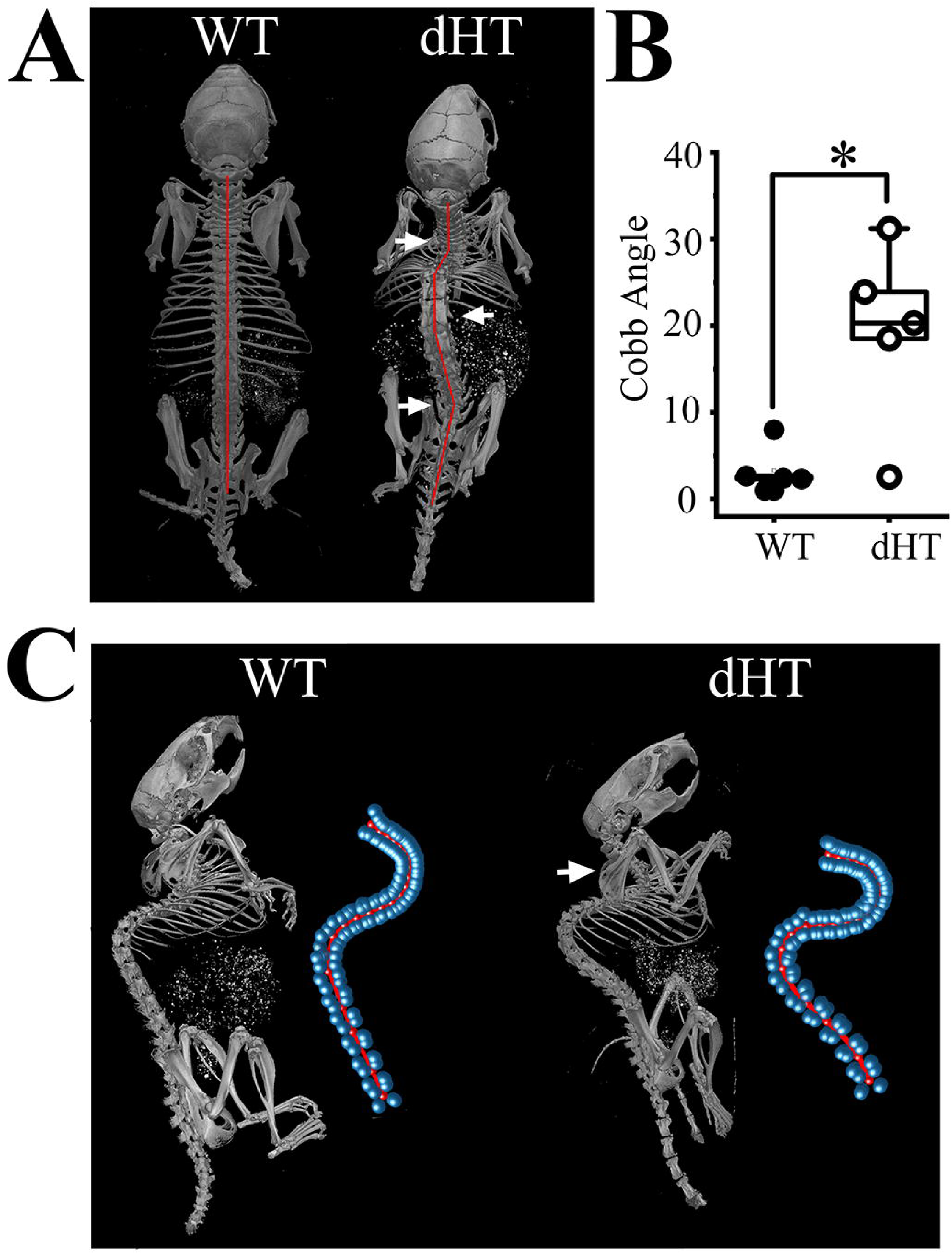
dHT mice show skeletal deformities, including scoliosis and kyphosis. Representative mCT imaging of skeletons from 12 weeks old WT and dHT mice **A.** Frontal view of the whole-body skeleton of a WT (left) and dHT (right) littermate. The red lines follow the spinal columns, the white arrows mark changes in the spinal alignment. **B.** Whisker plots of Cobb angles. Each symbol represents the Cobb value from a single mouse (WT n=6 and dHT n=6) (p<0.05 Mann Whitney test). **C.** Lateral views of whole- body skeletons from WT (left) and dHT (right) mice. A 3D reconstruction of the spine is shown on the right of each mouse. There are 8 blue spheres for each vertebra, and the red central line follows the path between the vertebras. Kyphosis is very pronounced in dHT mice (white arrow).

### Structural disorders of muscle spindle in dHT mice are paralleled by the severe depletion of RyR1 protein and significant alterations of the proteome in intrafusal muscle fibers

In the next section we investigated whether the presence of the mutant RyR1s is accompanied by reduction of RyR1 protein content in the muscle spindles, and if this is associated with changes of the proteomic signatures. High resolution confocal immunohistochemical analysis on cryosections of EDL and soleus muscles from dHT-Thy- EGFP mice (Fig.6A and 6B) did not reveal any major changes in the appearance of the nuclear clusters of the equatorial region of intrafusal muscle fibers as indicated by DAPI staining. Similar to what was observed in WT-Thy-EGFP mice, the RyR1 Ab preferentially stains the polar regions of the intrafusal fibers. RyR1 immunostaining of the intrafusal fibers from dHT shows the double row pattern, typical of the calcium release units of the junctional sarcoplasmic reticulum. Since immunohistochemistry is not a quantitative method, we compared RyR1 protein content in muscle spindles from soleus and EDLs isolated from dHT and WT mice by high resolution mass spectrometry. Fig.6C shows that the RyR1 protein content in intrafusal muscle fibers of dHT mice is decreased by 54.7% (p=0.0022) compared to that of WT mice.

**Figure 6:**
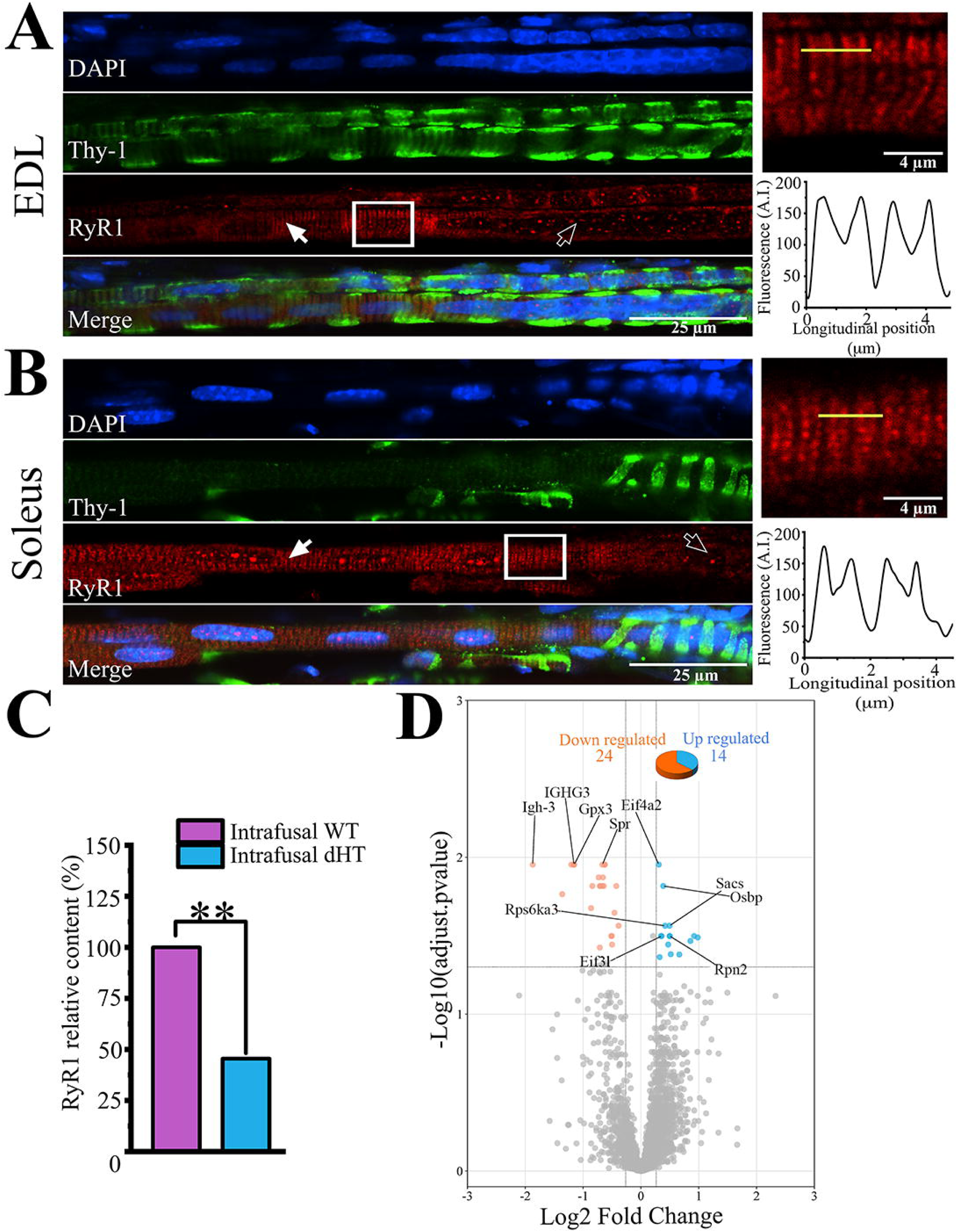
Immunohistochemical and proteomic analysis of intrafusal muscle fibers in dHT mice. A. and B. Subcellular localization of RyR1 protein in muscle spindle fibers from 12 weeks old dHT mice, Confocal images were acquired along the longitudinal axis of muscle spindle fibers from EDL (**A**) and soleus muscles (**B**). Images show the general structure of the muscle spindle. In the polar region (white arrows) the RyR1 positive immunofluorescence is highest. Fibers were fixed and processed as described in the Methods section, and stained with DAPI (blue), Thy-1 (green) and RyR1 (red). Images were acquired using a Nikon AxR confocal microscope equipped with a 40x Plan Apo VC Nikon objective (NA= 0.95). Scale bars = 50 µm. The two insets show a higher magnification of the boxed region in EDL and soleus RyR1 staining, where the double labelling pattern characteristic of RyR1 distribution is clearly visible. White magnification bar = 4 µm. The yellow bar shows the longitudinal position of the fluorescently labelled RyR1. The lower panels of the two insets show the longitudinal spatial fluorescence profiles over the transverse direction within the line. **C**. Mass spectrometry quantification shows that the RyR1 protein in intrafusal fibers from dHT mice is decreased by 54.7% compared to WT littermates. Results are the mean of muscle spindles from 5 mice per genotype. **D.** Volcano plot comparison of proteins in muscle spindles from WT versus dHT littermates. After filtering data stringently (keeping only proteins having ≥ 2peptides, exhibiting a fold change ≥0.20 (-0.321≤Log2FC≤0.263) and a q value ≤0.05) 24 proteins were down-regulated and 14 up-regulated in spindles from WT versus dHT, respectively.

As depicted in the volcano plot in Fig.6D, the protein composition of intrafusal fibers from dHT mice differs from that of WT mice with >30 proteins showing significant changes after filtering for those proteins showing ≥2 peptides, exhibiting a fold change ≥0.20 (- 0.321≤Log2FC≤0.263) and a q value ≤ 0.05. A total of 14 proteins were up-regulated and 24 proteins were down-regulated in intrafusal muscles from dHT vs WT; Table 2 shows the complete list of up-regulated and down-regulated proteins. The most affected pathways are those involved in signaling regulation, cellular amide biosynthesis and cellular organization that are significantly increased in dHT vs WT intrafusal fibers (Supplementary Fig. 7A). Whereas, pathways involved in cell metabolism, biosynthesis and peptide metabolism (Supplementary Fig. 7B) are significantly down-regulated in dHT intrafusal fibers. The proteins showing the greatest enrichment in intrafusal fibers from dHT vs WT mice are: (i) prolyl4-hydroxylase subunit α2 a subunit of a key enzyme involved in the proper three-dimensional folding of newly synthesized procollagen chains; (ii) Prepl prolyl endopeptidase a ptolyl endopeptidase-like enzyme; (iii) transferrin receptor (see Table 2). Of interest, Stromal interaction molecule 1 (or Stim1) a transmembrane protein mediating calcium influx after calcium store depletion, was increased 1.43-fold (q value= 0.041) in dHT vs WT intrafusal fibers. On the other hand the proteins showing the largest decrease in dHT intrafusal fibers are ß-IGH3 a TGF-ß inducible protein implicated in connecting matrix molecules to each other and facilitating cell-collagen interactions, NAD(P)H quinone dehydrogenase 1 a cytosolic FAD-binding enzyme involved in the reduction of quinones to hydroquinones and in the ubiquitin-independent p53 degradative pathway, ribosomal protein L28 (Table 2) (see https://massive.ucsd.edu/ProteoSAFe/private-dataset.jsp?task=09ff33af463e430f8e40ce18bf5cf1d2 for complete Mass Spectrometry datasets).

**Table 2:**
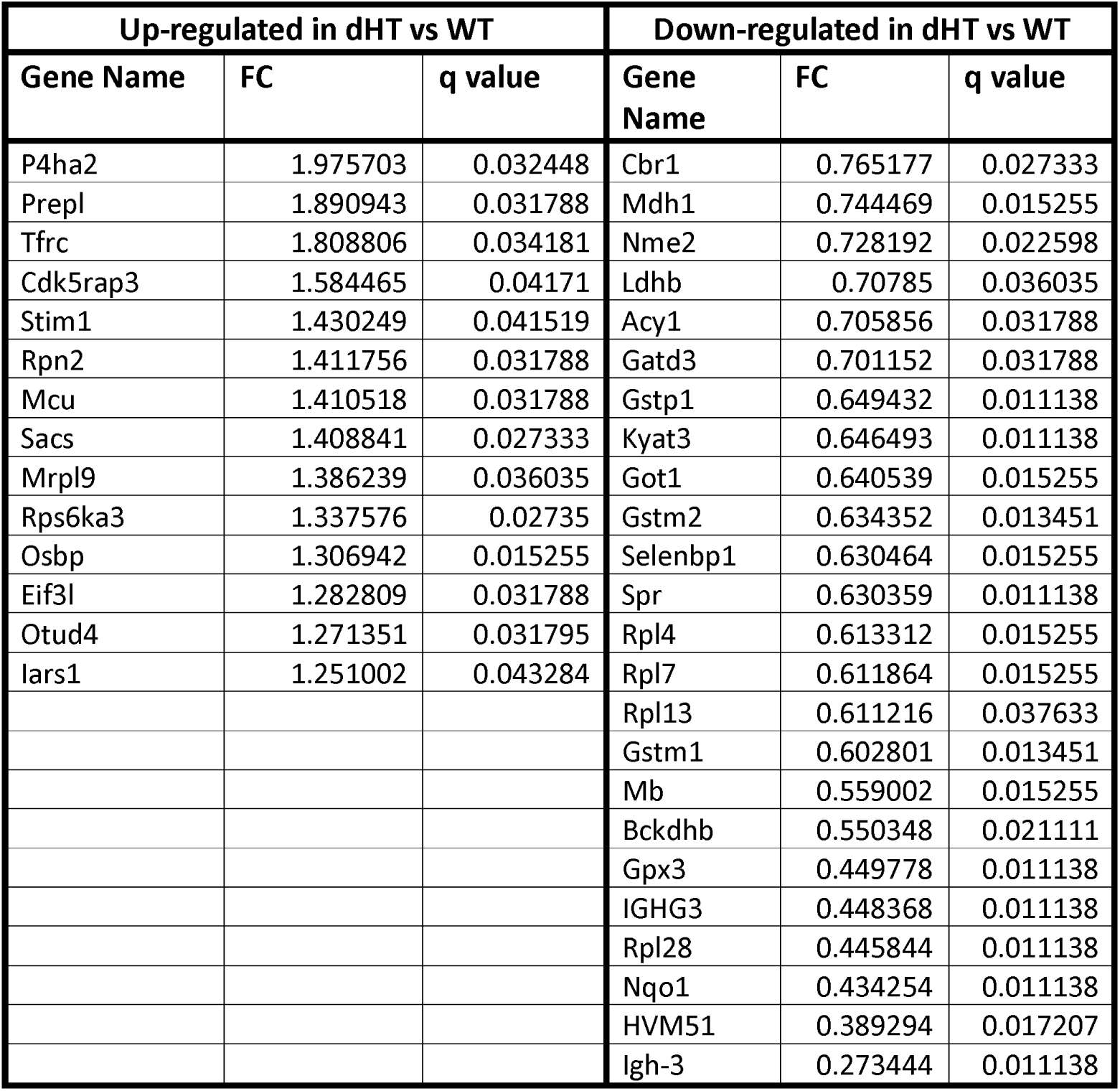
List of significantly up-regulated and down-regulated proteins in dHT vs WT intrafusal muscles.

## Discussion

In the present report we demonstrate that intrafusal muscles express the RyR1 protein and that *in vivo* parameters associated with proprioceptor functions are affected in dHT mice carrying one hypomorphic allele and one allele containing the p.A4329D missense RyR1 mutation. The observed changes in muscle spindle protein composition brought about by the presence of RyR1 mutations, causes a massive decrease of RyR1 expression in intrafusal fibers which may be consistent with the gait abnormalities observed in dHT mice and for the gross alterations of muscle spindle structure. If these alterations observed in dHT mice also occur in humans, our results could explain at least in part, the skeletal abnormalities, motor delays, joint contractures and gait abnormalities often observed in patients with *RYR1*-congenital myopathies.

### Muscle spindles show a unique protein composition

We used laser capture microdissection (51), to define the proteome of murine intrafusal muscle fibers. Our results demonstrate the validity of our experimental approach since, intrafusal muscles are endowed with a unique proteome compared to extrafusal fibers which are enriched in pathways involved in muscle contracture, sarcomere organization, oxidative phosphorylation and mitochondrial function. In particular they are enriched in proteins involved in extracellular matrix organization, collagen binding, chromatin assembly and filopodium assembly. Importantly, this study confirms and extends single nucleus transcriptomics analysis (43) and mouse masseter muscle spindle mass spectrometry data (44). Specifically, we confirm the presence of several proteins identified in murine muscle spindles including Padi2 identified in nuclear Bags (43) and within intrafusal fibers (44, 52). Deficiency of Padi2 leads to loss of bone mass by interfering with the transcription factor Runx2 (Cbfa1) an essential master of transcription of skeletogenesis (50, 53, 54).

By Mass Spectrometry Bornstein et al. identified a total of 40 proteins that are uniquely up-regulated in muscle spindles compared to the surrounding extrafusal masseter muscles (44). Here we identified more than 1500 up-regulated proteins, some of which are in common with those observed by Bornstein et al, including Tubb3, Vcan, Eln, Efmp1, Postn, ATP1a3, Hspa12a, Vat1l, Plp1, Myl4; Myl6b and Padi2. We were unable to confirm the enrichment of Slc2a1 encoding the Glut1 transporter, most likely because it is expressed in the outer-most portion of the capsule (44), an area that was not collected using laser capture micro-dissection. Of relevance, muscle spindles contain the RyR1 protein, albeit to a much lower extent than that present in extrafusal muscles.

### Alterations of the structure of the spindle in dHT mice

Muscle spindles are composed of intrafusal muscle fibres surrounded by a thin spindle-shaped capsule made up of specialized cells and connective tissue whose role is postulated as being a pressure sensory organ (55) and/or a metabolically-active diffusion barrier to the entrance of substances from the external milieu (37). The broadness of the gap between muscle spindles and extrafusal fibres is significantly increased in muscles of dHT mice, a finding that has also been reported to occur in other neuromuscular disorders and/or animal models. For example, in muscle spindles of patients with Duchenne muscular dystrophy the gap between muscle spindles and the extrafusal fibre region becomes significantly wider (33, 56), Similarly, this occurs also after denervation and during aging (57, 58). We did not find changes in gap thickness between muscle spindles and extrafusal fibres in Ex36 and Ho mice, thus the presence of *Ryr1* mutations alone is not sufficient to cause changes in the broadness of the gap between muscle spindle.

### Alteration of proprioceptive function in dHT mice

We are aware that the Balance beam and CatWalk tests don’t directly evaluate only muscle spindle function. Nevertheless, they do appraise proprioceptor parameters including motor coordination and balance and have been used for the early detection of motor deficits in mouse models of Huntington’s disease, in ALS, Parkinson’s disease, Pompe’s disease (59–62). CatWalk gait analysis revealed a decreased Base of Support in dHT. The alteration of proprioception can lead to a reduction in the base of support due to several reasons related to the body’s ability to perceive and respond to its position and movement in space. A reduction of the base of support was also reported in a murine model of Refsum disease (63) a rare metabolic disease accompanied by ataxia and skeletal abnormalities Gait abnormalities might be due to a variety of factors including ergogenic deficit of extrafusal fibre as well as to alteration of proprioceptive properties linked to muscle spindle disfunction. The gait and interlimb coordination disorders observed in dHT mice might be due a several factors including (i) deficit of muscle strength of extrafusal fibres, (ii) alteration of the contractility of intrafusal fibers upon γ motor neuron dependent stimulation, or (iii) the combination of both. Although we cannot discriminate between these different possibilities, our results suggest that the deficit of muscle strength of extrafusal fibres per se might not be the only factor that should be taken into consideration to explain gait and interlimb coordination abnormalities observed in dHT mice. This argument is consistent with the observation that the Ex36 mouse model displays no major changes of gait and interlimb coordination properties and yet it shows an approx. 20 % decrease of muscle strength and reduction of RyR1 protein content in extrafusal fibres (Supplementary Fig.2). The dHT mice exhibit massive reduction of RyR1 protein content in both intrafusal and extrafusal fibres, in the latter, the decrease of RyR1 protein is associated with a decrease of the amplitude of calcium transients and muscle strength (38). Based on previous (41) and present high resolution TMT mass spectrometry results, we calculated that the RyR1 tetrameric complex content in intrafusal fibres from dHT is 0.071 µmol/kg wet weight, a value which 45% of that present in WT intrafusal fibres. Thus, the massive reduction of RyR1 protein content in intrafusal fibres from dHT mice would cause a decrease of intrafusal contractility mediated by the RyR1 calcium release occurring during the γ motor neuron stimulation. This could in turn impair the continuous adjustment of muscle spindle activity during normal locomotor activity. If this is the case then the deficit of the muscle spindle adjustment occurring in dHT mice is a potential component that should be taken into account when considering the deficit of its proprioceptive system leading to a variety of effects including scoliosis. In this context, it should be mentioned that patients with some recessive *RYR1* mutations often show scoliosis and joint contractures from birth. Indeed, in a recent retrospective study of 69 patients with *RYR1* mutations, Sarkozy et al. showed that 71% patients with AR mutations presented signs at birth, 19% had contractures including contractures of the neck, shoulder, elbow, wrists, thumb, hip, knee and ankles at birth, 23% had scoliosis and 29% had spinal rigidity (19). Similarly, Ambourgey et al. reported that 100% of the patients with recessive *RYR1* mutations presented symptoms and of these, 50% had scoliosis and 71% contractures (4) which appeared at birth, before the development major motor milestones such as deambulation.

### The proteomic composition of muscle spindles from dHT mice is altered

Since the expression of RyR1p.Q1970fsX16+p.A4329D mutations causes significant changes in the protein composition of extrafusal muscles (41), we were interested in investigating the proteome of intrafusal muscles of dHT mice. The proteome of dHT intrafusal fibres shows that a total of 14 proteins are up-regulated and 24 proteins are down-regulated in dHT compared to WT mice. Interestingly, 21% (5/24) of the down- regulated proteins can either bind to, or are involved in, glutathione metabolism (carbonyl reductase 1, glutathione S-transferase pi 1, glutathione S-transferase Mu 1 and 2 and glutathione peroxidase 3) and 21% are ribosomal constituents. Down-regulation of proteins involved in glutathione metabolism could lead to protein carbonylation, oxidative stress and mitochondrial dysfunction affecting the function of many proteins, including the RyR1 a major target of oxidative stress (64). Ribosomal proteins are principally involved with the stabilization of the ribosomal subunits, rRNA processing, RNA folding etc however, they also possess extra-ribosomal functions including regulation of cell growth, proliferation, DNA repair and more. Thus, a decrease in large subunit ribosomal proteins content may be linked to altered mRNA processing and/or protein synthesis within the muscle spindles. In the context of RYR1-related congenital myopathies the finding that RPL13 is significantly down-regulated in dHT muscle spindles may be of relevance. Indeed, mutations in RPL13, have been identified in patients with Spondyloepimetaphyseal dysplasia, a disorder characterized by short stature, scoliosis and kyphosis, hip dislocation, equinovarus foot (club foot) and other orthopaedic joint conditions (65), a large part of these symptoms are shared by patients carrying recessive *RYR1* mutations.

In the case of up-regulated genes, the protein showing the largest increase in intrafusal of dHT versus WT is collagen prolyl 4 hydroxylase type 2. Collagen prolyl 4 hydroxylase are capable of hydroxylating the majority of proline sites in type 1 collagen promoting the formation of the triple helix. Thus, the increased content of collagen prolyl 4- hydroxylase may be a down-stream effect linked to the abnormal content of collagen in intrafusal muscles fibres from dHT mice.

### Conclusions

In this study we provide unambiguous evidence for the expression of RyR1 in intrafusal muscle fibers and demonstrate that the massive RyR1 reduction associated with recessive *Ryr1* mutations together with the presence of a missense mutation, impact gait and interlimb coordination and cause significant dysregulation of protein expression. These effects may be involved in the skeleton abnormalities observed in patients affected by recessive *RYR1* mutations. A limitation of our study is that we cannot determined whether the altered function of the RyR1 calcium channel is the only cause of the muscle spindle changes or whether these are secondary to changes of protein expression brought about by the presence of the mutations.

## Materials and Methods

### Mice and compliance with ethical standards

All experiments were conducted on 11-12 weeks old male mice unless otherwise stated. Experimental procedures were approved by the Cantonal Veterinary Authority of Basel Stadt (BS Kantonales Veterinäramt Permit numbers 1728 and 2115. All experiments were performed in accordance with relevant guidelines and regulations. The following mouse models were used: dHT carrying the compound heterozygous RyR1p.Q1970fsX+p.A4329D mutations, Ex36 carrying the RyR1p.Q1970fsX mutation, Ho carrying the homozygous RyR1.pF4976L mutation, their WT littermates, as well as B6-Cg-Tg (Thy1-YFP) mice from Jackson laboratories. The latter were also bred with dHT and Ex36 transgenic mice for confocal immunohistochemical analysis on muscle spindles

### Mouse Genotyping

PCR amplification of *Ryr1* exons 36, 91 and 104 was performed as previously described (38, 40) on genomic DNA of WT, Ex 36, RyR1Q1970fsX16+A4329D and Ho (RyR1F4976L) transgenic mice. B6-Cg-Tg (Thy1-YFP) genotyping was carried out as described (https://www.jax.org/Protocol?stockNumber=003709&protocolID=32064) using Probe Based qPCR and an Internal Positive Control (IPC) as a reference, except that the qPCR kit from Promega (catalog N° A6102) was used. Each primer set was used in combination with a probe for fluorescence quantification during the qPCR amplification reaction. The complete list of primers used for genotyping is given in Supplementary Table 3.

### Determination of spindle diameter by hematoxylin and eosin staining

Soleus muscles were isolated from 12 weeks old male mice, embedded in PolyFreeze (Sigma, catalog N°P0091), deep-frozen in 2-methylbutane (Sigma, catalog N° M32631) and stored at −80°C. Muscles were then cut in 10 µm thick sections using a Leica Cryostat (CM1950) and stored at -80°C. Samples were then stained with hematoxylin and eosin (H&E) using a commercial staining kit (Abcam, catalog N°245880). Sections were visualized using a 40x Plan Apo λ objective (N.A.=0.95) using an Eclipse Ti2 Nikon widefield microscope. The diameter of muscle spindles (distance of parallel tangents at opposing borders of the fiber) was calculated from the central region of the muscle spindle (equatorial region) which was recognized after scrolling through the individual images of the stack (45). Images were analyzed using the Fiji software (National Institutes of Health, Bethesda, Maryland, USA).

### Immunohistochemical analysis of muscle spindles

Twelve weeks old male or female mice were deeply anesthetized by an intraperitoneal injection of pentobarbital and perfused with 4% PFA for fixation. Fixed EDL and soleus muscles were isolated and embedded in PolyFreeze medium (Sigma, catalog N°P0091), frozen in cold hexane and stored at -80°C until used. Longitudinal 25 µm sections were obtained using a Leica Cryostat (CM1950) and processed as described (62). For staining, sections were first rehydrated in PBS for 10 min. Antigen retrieval was performed by steaming in the immunohistochemistry antigen retrieval buffer (Thermo Fisher Scientific, catalog N°00- 4955-58). Blocking was performed using a blocking solution (western blocking reagent Roche catalog N°11921673001) containing 0.15% Triton X-100 and 1% BSA in 300 mM Glycine for 1 hour at room temperature. Subsequently, sections were incubated overnight at 4°C with the following primary antibodies: rabbit D4E1 monoclonal anti-RyR1 (Cell Signaling Technology, catalog N°8153; 1:150 dilution in Phosphate Buffer Saline, PBS), chicken anti-GFP to visualize YFP-Thy1 (Abcam catalog N° ab13970; 1:500 in 1% BSA in PBS). The next morning samples were washed at room temperature 3 times 10 min each with 0.1% Tween in PBS (PBS-T) and then incubated for 1 hour at room temperature with the following secondary antibodies: goat anti-rabbit Alexa Fluor 647 (Thermo Fisher Scientific, catalog N° A21245; 1:1000 dilution in PBS-T) and goat anti-chicken Alexa Fluor 488 (Thermo Fisher Scientific, catalog N° A11039; 1:1000 dilution in PBS-T). Samples were washed 2x 10 min each with PBS-T and 1x for 10 min with PBS and then incubated with 4′,6-Diamidine-2′-phenylindole dihydrochloride (DAPI) (Sigma catalog N° D9542; 1:500 in PBS) for 15 min at room temperature. Finally, samples were washed 2x 10 min each with PBS and 1x for 10 min, mounted in Fluoromount G (Thermo Fisher Scientific, catalog N°00-4958-02) and visualized using a confocal Nikon AxR microscope equipped with a 40x Plan APO objective (N.A. =095).

### Laser capture microdissection (LCM) and mass spectrometry

Soleus and EDL muscles isolated from 11-13 weeks old male mice were embedded in OCT (Sigma), deep-frozen in 2-methylbutane (Sigma), cut in 25 µm thick sections as described above and stored at -80°C. Spindles identified in consecutive cross-sections were isolated by laser capture microdissection as described below, using a PALM Robot-microbeam system (P.A.L.M. Microlaser Technologies AG) equipped with the PALM RoboSoftware version 1.2 (P.A.L.M. Microlaser Technologies AG) and mounted on a Zeiss Axiovert 200 M microscope (Carl Zeiss, Oberkochen, Germany). Muscle spindles identified within a muscle section, were encircled in an “region of interest” and dissected out of the tissue using the cutting and catapulting function, RoboLPC. The catapulted material was collected in the cap of a 200-µL Thermo-Tube (ABgene, Epsom, UK) containing 20 µL lysis buffer (5% SDS; 50 mM Triethylammonium bicarbonate, TEAB). Twenty-five microdissected sections from the polar area of the muscle spindle were isolated per mouse.

For mass spectrometry, 10 mM tris 2-carboxyethyl phosphine (TCEP) was added to each sample, followed by heating to 95 °C for 10 min and sonication (Bioruptor, 10 cycles, 30 s on/off, Diagenode, Belgium). After equilibrating to room temperature, iodoacetamide (IAA) was added and the samples were then incubated at room temperature (protected from light) for 30 min. Proteins were then purified and digested following the modified SP3 protocol (66). In brief, Speed Beads^TM^ (#45152105050250 and #65152105050250, GE Healthcare) were mixed 1:1, rinsed with water and diluted to the 8 μg/µL stock solution. Samples were adjusted to the final volume of 25 µL and 5 µL of the beads stock solution was added to the samples. Proteins were bound to the beads by addition of 30 µL 100% acetonitrile to the samples (to the final concentration of 50%), which were then incubated for 8 min at RT with a gentle agitation (200 rpm). After, samples were placed on a magnetic rack and incubated for 5 minutes. Supernatants were removed and discarded. The beads were washed twice with 200 µL of 70 % (v/v) ethanol and once with 200 µL of 100% acetonitrile. Samples were placed off the magnetic rack and 10 µL of digestion mix (5 ng/µL of trypsin in 0.02% n-Dodecyl-Beta-Maltoside, 50 mM triethylammonium bicarbonate) was added to them. Digestion was allowed to proceed for 12h at 37°C. After digestion samples were placed back on the magnetic and incubated for 5 minutes. Supernatants containing peptides were collected and dried under vacuum.

Dried peptides were dissolved in 10 µl of 0.1% formic acid and 4 µl of the sample was subjected to LC–MS/MS analysis using an Orbitrap Eclipse Tribrid Mass Spectrometer fitted with an Ultimate 3000 nano-LC (both Thermo Fisher Scientific) and a custom-made column heater set to 60°C using block randomization.

Peptides were resolved using a RP-HPLC column (75μm × 30cm) packed in-house with C18 resin (ReproSil-Pur C18–AQ, 1.9 μm resin; Dr. Maisch GmbH) at a flow rate of 0.3 μLmin-1. The following gradient was used for peptide separation: from 2% B to 12% B over 5 min to 30% B over 40 min to 50 % B over 15 min to 95% B over 2 min followed by 11 min at 95% B. Buffer A was 0.1% formic acid in water and buffer B was 80% acetonitrile, 0.1% formic acid in water. For the data acquisition, a FAIMS device attached and set to constant CV voltage of -45. The mass spectrometer was operated in DIA mode with a cycle time of 3s. MS1 scans were acquired in the Orbitrap in centroid mode at a resolution of 60,000 FWHM (at 200 m/z), a scan range from 400 to 1000 m/z, normalized AGC target set to 250 % and maximum ion injection time mode set to 50 ms. MS2 scans were acquired in the Orbitrap in centroid mode at a resolution of 120,000 FWHM (at 200 m/z), precursor mass range of 400 to 900, quadrupole isolation window of 56 m/z with 1 m/z window overlap, normalized AGC target set to 2000 % and a maximum injection time of 246 ms. Peptides were fragmented by HCD (Higher-energy collisional dissociation) with collision energy set to 27 % and one microscan was acquired for each spectrum.

The acquired raw-files were searched by directDIA against the murine UniProt protein database (version Feb 2022) and commonly observed contaminants) by the SpectroNaut software (Biognosys, version 18.6.231227.55695) using default settings. The search criteria were set as follows: full tryptic specificity was required (cleavage after lysine or arginine residues unless followed by proline), 2 missed cleavages were allowed, carbamidomethylation (C) was set as fixed modification and oxidation (M) and N-terminal acetylation as a variable modification. The false identification rate was set to 1%. The search results were exported from SpectroNaut and protein abundances were statistically analyzed using MSstats (v.4.4.1) (67). Genes showing a ≥2.0-fold change in protein expression (p ≤ 0.05) were analyzed by DAVID functional annotation to produce gene clusters (≥2 genes/cluster). GO terms corresponding to biological process (GOTERM_BP_FAT and KEGG_PATHWAY) were extracted and are plotted with the numbers of genes (as a percentage of the total) for each term. GO terms with <2% of the total genes were not plotted unless significantly enriched (Benjamini ≤0.05).

### *In vivo* proprioceptor functional tests using the Beam Walk and Catwalk tests

All behavioral tests and analyses were conducted by an experimenter blinded to the mouse genotype. Mice were investigated for 10 weeks, starting from the age of 6 weeks. The first week was used as acclimatization to the experimental device. In the case of the Beam walk test, balance was tested on a 1 m long, 1 cm wide beam suspended on two poles 50 cm above a table top. Food was placed in a house-like goal structure at the endpoint to attract the mouse. A video camera on a tripod was used to record the walk and help the experimenter assign the score for each mouse. Scores were given on a scale of 0 – 3, zero representing absence of any pathological phenotype, 3 representing a compromised phenotype (see Supplementary Figure 4A for score details). Scores were assigned as follows: 0 in the case of unaltered balance during the beam crossing; 1 if a mouse misplaced its foot on the beam but walked in a coordinated manner; 2 in the case of unbalanced behavior, continuous foot slipping or dragging of posterior legs; 3 in the case of falls and/or inability to move despite encouragement. Beam walk measurements were conducted four times per week and the last two measurements were used for the statistical analysis, since these were the ones when mice were the most acquainted to the experimental procedure. CatWalk XT system (v10.0.408; Noldus, USA) was used to assess qualitative and quantitative differences between WT and dHT age matched male mice following published experimental protocols (49). Supplementary Fig.5 outlines how parameters were assessed. Prior to testing, mice were acclimatized to the dark room and the CatWalk XT instrument over three days for 1 h per day with the illuminated surface turned on. The CatWalk system automatically recorded videos of the mice walking the entire length of the walkway. Experimental sessions typically lasted 5-10 min, and three compliant runs were recorded per mouse. Successful runs were established as a continuous and straight walk, without interruptions or head turning to the sides. Camera gains was set to: 20dB, detection threshold of 0.1 with red ceiling light of 17.7V and green walkway of 16.5V. Experiments were conducted at the same time of the day and a minimum of three successful runs was acquired. The CatWalk system determined the compliance (60%) according to the run’s duration and speed variation. Paw positions were verified manually after acquisition. Videos were classified and data were analyzed using custom software and the results of three compliant runs per mouse were averaged and plotted. CatWalk data sets are presented in Supplementary Figure 4C and 4D and in Supplementary Table 2.

### X-ray microtomography

Twelve-week-old male mice were sacrificed by intravenous injection of pentobarbital and immersed overnight in PBS containing 4% PFA. The whole mice were examined by X-ray microtomography using a SKYSCAN 1275 (Bruker, Kontich, Belgium) equipped with a 100 kV / 10 W microfocus X-ray source. All scans were performed with an accelerating voltage of 80 kV and a beam current of 125 µA. To increase the mean energy of the X-ray spectrum, a 0.8 mm Al filter was used. The effective pixel size was set to 20 μm and the exposure time to 0.63 s per rotation angle. 1440 equiangular projections were acquired over the angular range of 360°, resulting in an acquisition time of approximately 1.5 h per scan. Due to the size of the mouse, five height steps were recorded. The total acquisition time for the entire mouse was therefore around 7.5 hours. The projections were reconstructed with a cone beam filtered back projection algorithm using the manufacturer’s NRecon (version 1.7.4.6) software. VGStudio MAX 2.1 software (Volume Graphics GmbH, Heidelberg, Germany) was used for visualization and extraction of vertebral coordinates. A digital 3D reconstruction of the spine was performed to examine scoliosis and kyphosis. The Cobb angles were measured with Fiji (doi:10.1038/nmeth.2019).

For Kyphosis a Permutation statistical test, cubic splines were fit to the 3D point clouds of each mouse in the dataset (https://zenodo.org/doi/10.5281/zenodo.12721977). These smooth curves were constrained to be flat (zero curvature) at their extremes, corresponding to the points that would lead to the mouse 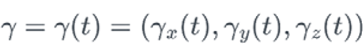 tail and head connections. That was made to focus on the inner part of the annotated spine columns and avoid spurious curvature at the extremes.

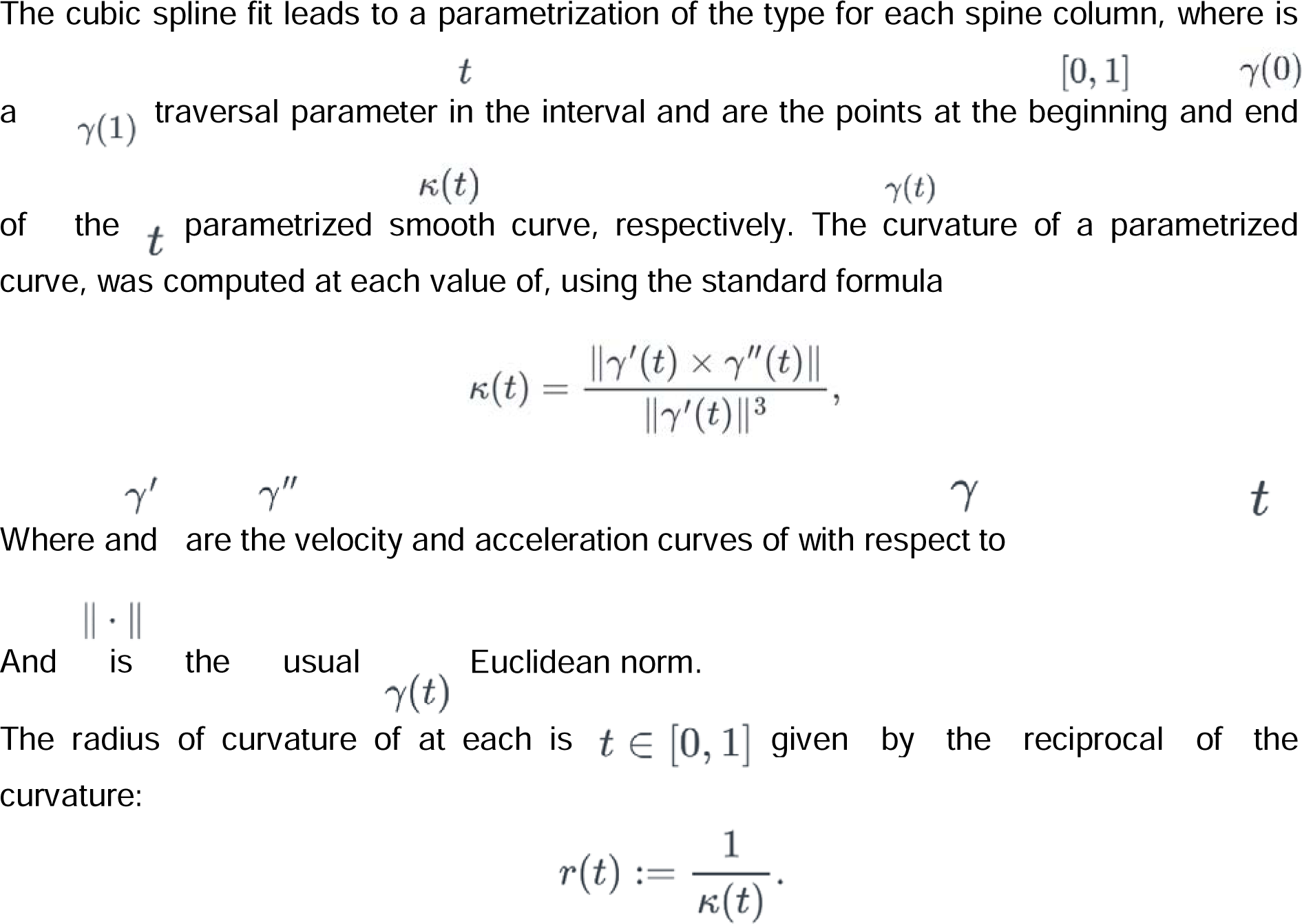

We compared populations of mice via summaries of the curvatures of the smooth curves fitted to their spine column point clouds. code used for the analysis is available at https://gitlab.com/ceda-unibas/spine-curvatures.

### Statistical analysis

Statistical analysis was performed using the Origin (Pro), Version 2019 software (OriginLab Corporation, Northampton, MA, USA.), using either Mann– Whitney test P values smaller or equal to 0.05 were considered significant.

## List of Abbreviations

CCD, central core disease; CRU, calcium release units; DHPR, dihydropyridine receptor; dHT, transgenic mice knocked in for compound heterozygous mutations, ECC, excitation-contraction coupling; EDL, *extensor digitorum longus*; FDB, *flexor digitorum brevis*; MmD, multiminicore disease; SR, sarcoplasmic reticulum; TT, transverse tubules; WT, wild type mice.

## Author contributions

Conceptualization: FZ and ST

Methodology: AR, CH, FZ, GS, HM, KB, RP, SB, ST,

Investigation: AR, FZ, GS, HM, KB, SB, ST Funding acquisition: ST, FZ

Project administration: ST, FZ Supervision: ST, FZ, AR

Writing – original draft: AR, FZ, ST

Writing – review & editing: FZ, ST, AR, SB, MF, RP, LP, FZ, FP

## Acknowledgements

We would like to thank Prof. Stephan Kröger for constructive criticism and suggestions.

## Data and materials availability

All data, code, and materials used in the analysis are available in some form to any researcher for purposes of reproducing or extending the analysis. There aren’t any restrictions on materials, such as materials transfer agreements (MTAs). All data are available in the main text or the supplementary materials. The Mass spectrometry data has been deposited in the ProteomeXchange Consortium via the ProteoSAFe repository (http://massive.ucsd.edu/ProteoSAFe) with the following access number PXD054222 (https://massive.ucsd.edu/ProteoSAFe/private-dataset.jsp?task=09ff33af463e430f8e40ce18bf5cf1d2).

The complete dataset of skeleton analysis can be found at: https://zenodo.org/records/12724636.

