## Supplementary Tables 1, 2 Supplementary Figures 1-7 for "Massive reduction of RyR1 in muscle spindles of mice carrying recessive *Ryr1* mutations alters proprioception and causes scoliosis"

**Supplementary Table 1:** Enriched proteins (values are intrafusal vs extrafusal muscle fibers) identified in the current study as well as previously reported (Ref 43 and 44)

| **Gene Name** | **Fold change** | **Q value** | **Location** | **Reference** |
| --- | --- | --- | --- | --- |
| *Synm* | 0.523 | 0.0049 | Chain2 | 43 |
| *Mylk2* | 0.545 | 0.00079 | Chain2+Bag+spdNMJ | 43 |
| *Tnnt3* | 0.583 | 0.011 | Chain1, Chain2+Bag+spdNMJ+spdMTJ | 43 |
| *Myom1* | 0.617 | 0.00037 | Chain1, Chain2+Bag+spdNMJ | 43 |
| *Ache* | 1.676 | 0.017 | Bag+spdNMJ | 43 |
| *Col3a1* | 2.064 | 0.001 | Chain+Bag+spdMTJ | 43 |
| *Col6a1* | 2.299 | 1.11x10^-6^ | spdMTJ | 43 |
| *Hmgb1* | 2.537 | 8.71x10^-8^ | Chain1, Chain2+Bag+spdNMJ | 43 |
| *Myl6b* | 2.320 | 0.00027 | Intrafusal fiber | 44 |
| *Vat1l* | 3.119 | 0.012 | Neuron | 44 |
| *Eln* | 3.610 | 1.0x10^-8^ | Capsule | 44 |
| *Igfn1* | 4.228 | 0.00026 | Chain1, Chain2+Bag+spdNMJ+spdMTJ | 43 |
| *Tubb3* | 4.594 | 1.53x10^-7^ | Neuron | 44 |
| *Vcan* | 5.291 | 0.00087 | Capsule | 44 |
| *Efemp1* | 6.485 | 9.74x10^-9^ | Capsule | 44 |
| *Plp1* | 6.570 | 5.74x10^-8^ | Neuron | 44 |
| *Thy1* | 6.624 | 1.58x10^-9^ | Neuron projections including the group Ia and group II sensory afferent endings in the equatorial region of intrafusal muscle fibers | 32 |
| *Myh6* | 7.029 | 1.47x10^-5^ | Bag | 44 |
| *ATP1a3* | 10.482 | 4.72x10^-10^ | Neuron | 44 |
| *Hspa12a* | 11.194 | 4.58x10^-12^ | Neuron | 44 |
| *Myl4* | 11.700 | 3.78x10^-5^ | Intrafusal fiber | 44 |
| *Padi2* | 11.876 | 2.69x10^-11^ | Intrafusal fiber | 44 |
| *Postn* | 18.099 | 3.2x10^-9^ | Capsule | 44 |

**Supplementary Table 2**: Analysis of Catwalk parameters in WT, dHT and Ex36 mice. All values are expressed as % of the values obtained in WT mice.

| **Parameter** | **Paw** | **WT (n=12)** | **dHT (n=7)** | **Ex36 (n=6)** |
| --- | --- | --- | --- | --- |
| Stand | RH | 100.0±18.4 | 119.4±26.7 | 134.9±29.3*  (p=0.026) |
|  | LH | 100.0±21.8 | 124.1±24.7*  (p=0.026) | 105.8±31.6 |
| Stride Length Front  Paws | RF | 100.0±7.6 | 81.2±13.3**  (p=0.0089) | 92.9±13.6 |
|  | LF | 100.0±7.7 | 81.1±13.2**  (p=0.0071) | 97.2±13.3 |
| Stride Length Hind  Paws | RH | 100.0±11.9 | 79.6±11.5**  (p=0.003) | 87.9±14.6 |
|  | LH | 100.0±11.5 | 83.8±15.0*  (p=0.023) | 93.4±15.6 |
| Duty Cycle | RH | 100.0±16.6 | 123.2±11.7**  (p=0.0031) | 118.3±22.3 |
|  | LH | 100.0±16.5 | 117.6±12.9*  (p=0.039) | 107.9±30.0 |
| BOS | FP | 100.0±12.6 | 81.2±7.2*  (p=0.012) | 102.2±23.3 |
|  | HP | 100.0±9.2 | 83.4±6.2*  (p=0.0027) | 107.2±10.7 |
| Min Intensity | RH | 100.0±14.3 | 95.1±4.8 | 104.7±24.3 |
|  | LH | 100.0±12.3 | 88.2±5.7*  (p=0.017) | 102.4±22.7 |
| Standing on three | NA | 100.0±57.1 | 269.2±10.1**  (p=0.0077) | 177.0±111.6 |
| Print area | RH | 100.0±17.1 | 93.5±20.1 | 109.8±21.5 |
|  | LH | 100.0±16.6 | 80.1±106.4**  (p=0.0032) | 87.3±35.4 |
| Initial Dual Stance | RH | 100.0±99.9 | 283.3±128.5*  (p=0.011) | 131.3±104.9 |
|  | LH | 100.0±64.0 | 349.1±266.5*  (p=0.010) | 91.3±87.3 |
| Terminal Dual Stance | RH | 100.0±42.8 | 259.6±200.8*  (p=0.041) | 84.9±64.2 |
|  | LH | 100.0±106.2 | 349.7±140.5*  (p=0.010) | 105.22±78.9 |

**Supplementary Table 3:** Sequence of primers used for genotyping

| **Gene amplification details** | **Primer sequence** | |
| --- | --- | --- |
|  | **Forward** | **Reverse** |
| *Ryr1* Ex36 | 5’-TGC TGG CTT CAG AGT GAT CG-3’ | 5’-CGA GGG AAG TTG AGG TTG GG-3’ |
| *Ryr1* Ex91 | 5’-GAG ATG TTC GTG AGT TTC TGC GAG G-3’ | 5’-TGA GGG TTG TTC TTG GTG TAT TTG G-3’ |
| *Ryr1* Ex104 | 5’-GAC CAA CAA GAG CAA GTG AAG G -3’ | 5’-GTC TAA ACC CTA CCC ACA CG-3’ |
| Thy1-GFP | 5’-CAC AGA ATC CAA GTC GGA ACT-3’ | 5’-AAC AGC TCC TCG CCC TTG-3’ |
| IPC Thy1-GFP | 5’-CAC GTG GGC TCC AGC ATT-3’ | 5’-TCA CCA GTC ATT TCT GCC TTT G-3’ |
| TG_Thy1_probe | 5’-CAC CTA GAG GAT CTC GAG GGA TCC-3’  *5’ modification*: FAM  *3’ modification*: TAMRA |  |
| IPC_Thy1_probe | 5’-CCA ATG GTC GGG CAC TGC TCA A-3’  *5’ modification*: HEX  *3’ modification*: TAMRA |  |

**Supplementary** **figures**

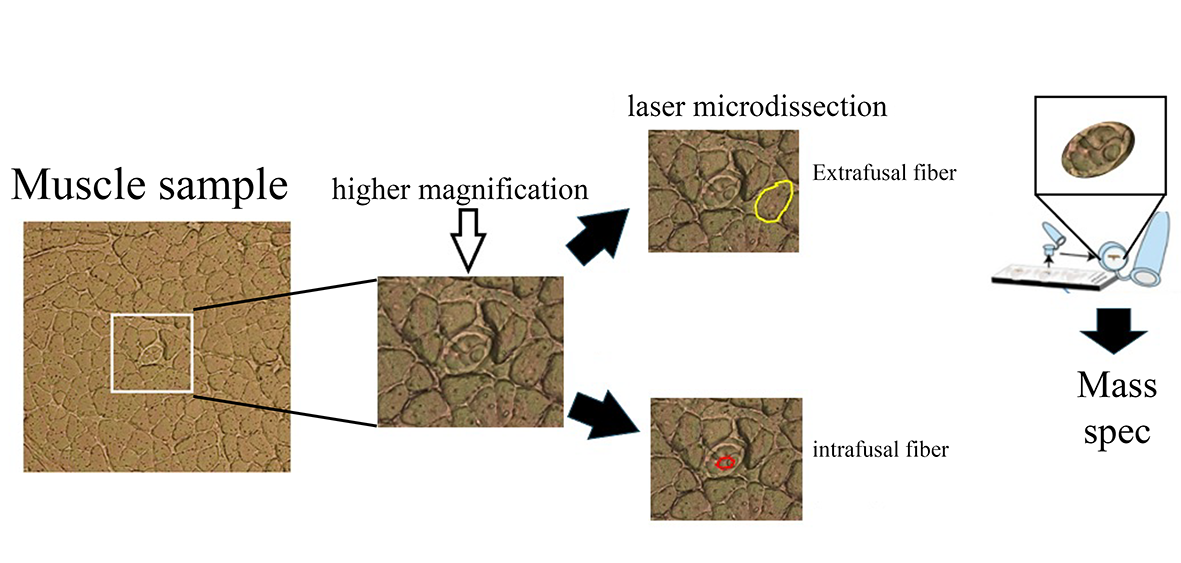

**Supplementary Figure 1:** Schematic representation of the experimental approach used to compare the protein composition of intrafusal muscle fibers and extrafusal muscle fibers from WT mice. Area of interest identified within a muscle section were collected by laser capture microdissection (LCM) and subsequently processed for quantitative proteomic analysis as described in the Methods section.

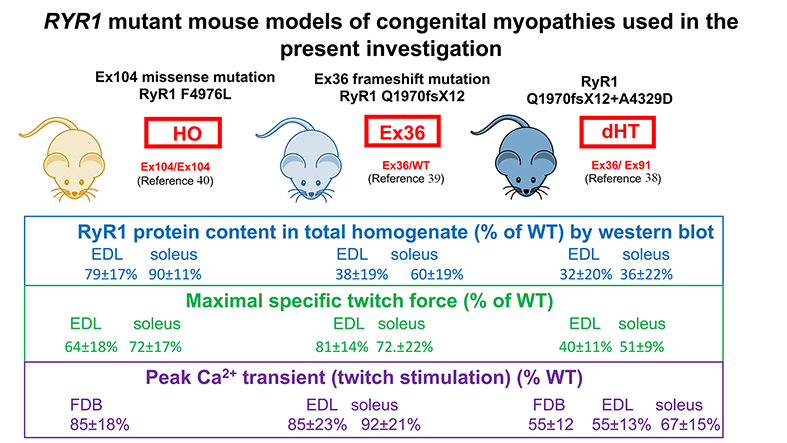

**Supplementary Figure 2:** Phenotypic characteristics of the transgenic mouse models used in the present investigation.

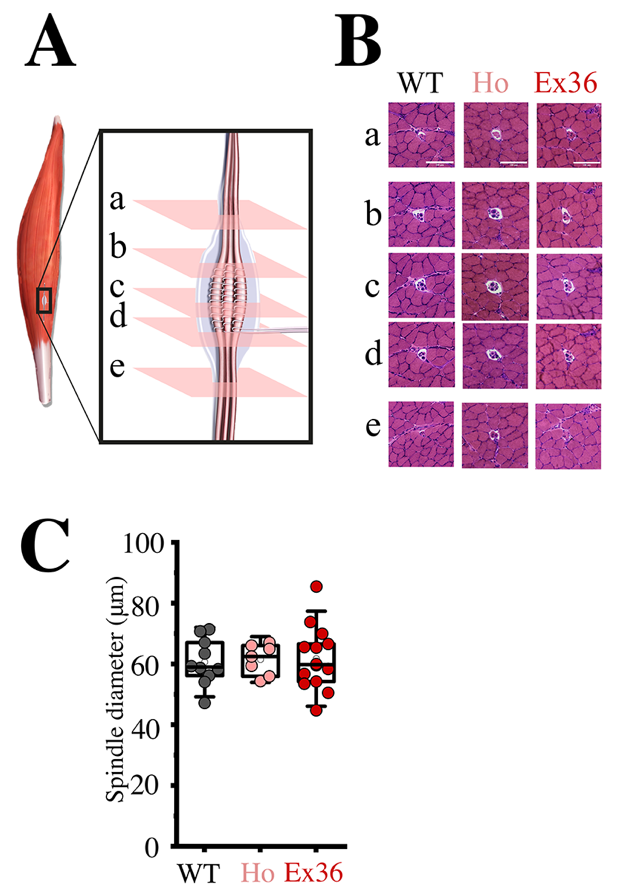

**Supplementary Figure 3: Ex36 and Ho mice do not exhibit gross alterations of muscle spindle morphology. A.** Schematic representation of a skeletal muscle showing the location of a muscle spindle. Skeletal muscles contain several longitudinally oriented muscle spindles ubiquitously distributed in the interior of the muscle belly. They are composed of intrafusal fibers (longitudinally oriented pink fibers) surrounded by a capsule (represented as a light blue matrix). Transverse planes (“a”, “b”, “c”, “d” and “e”) were made through the muscle spindles at different levels, “a” and “e” are located in the polar regions, “b”, “c” and “d” are located in the central region. **B.** H&E staining of soleus muscles sections containing intrafusal and extrafusal fibers from wild type (WT), Ex36 and Ho mice. Ten µm transversal muscle sections were made until the polar region was identified. White bar in panels “a” = 100 µm. Images were acquired with an Eclipse Ti2 Nikon widefield microscope. **C.** Whisker plots of muscle spindle diameter. The diameter was calculated in the equatorial section of the muscle spindle represented as region “c” in panels A and B. The spindle diameter (distance of parallel tangents at opposing borders of the fiber was calculated using ImageJ. Each symbol represents results obtained from a single spindle (11 spindles from n=6 WT mice; 7 spindles from Ho mice n=4 mice and 13 spindles from Ex 36 mice n=6 mice).

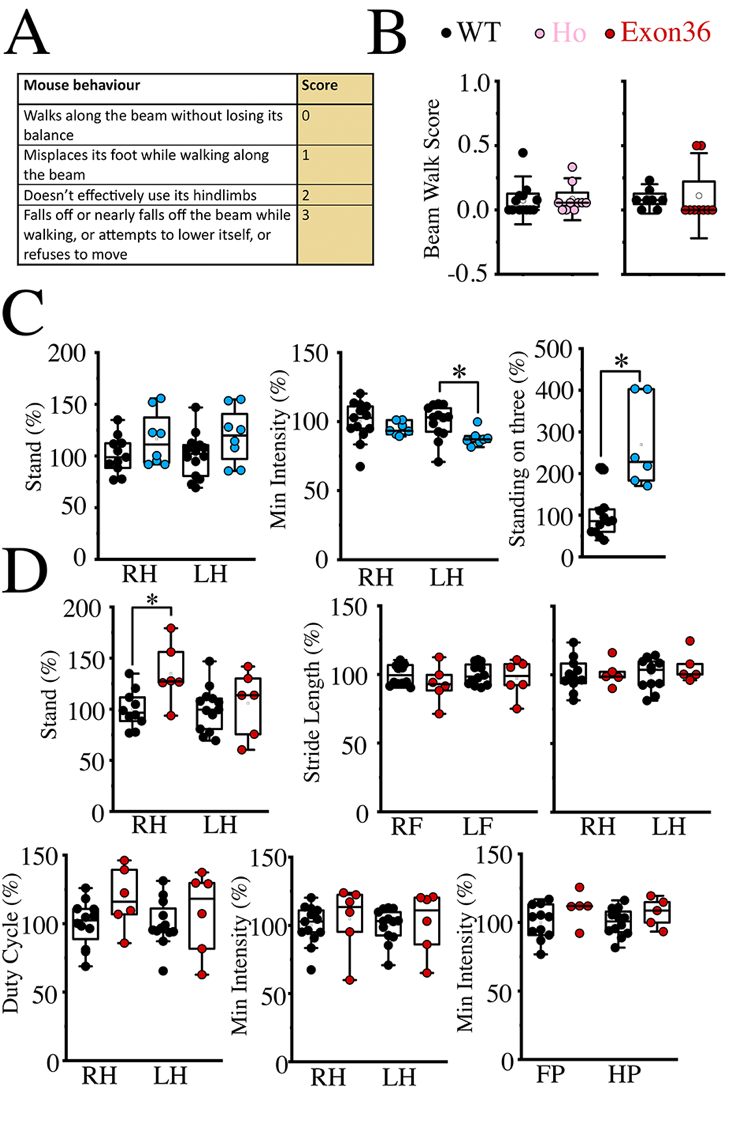

**Supplementary Figure 4:** **Ex36 and Ho mice do not show alterations in proprioceptor function. A.** Beam walk scoring parameters**. B.** The beam walk score tests of WT mice, Ex36 and Ho mice are similar; WT (n= 8), Ex36 (n=13), Ho mice (n=10). **C.** Catwalk gait analysis test performed using the CatWalk XT system, showing the Stand, min Intensity and Standing on three values in WT and dHT mice. **D.** Catwalk gait analysis test performed using the CatWalk XT system, showing the Stand, stride length, standing on three, Duty cycle and Min intensity of Right and Left Hindlimb and Front Paws values in WT vs Ex36 mice.

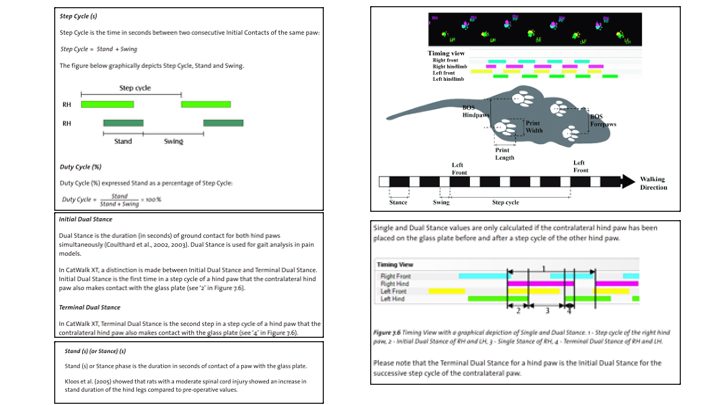

**Supplementary Figure 5:** Main gait parameters and their explanation, measured by the CatWalk system.

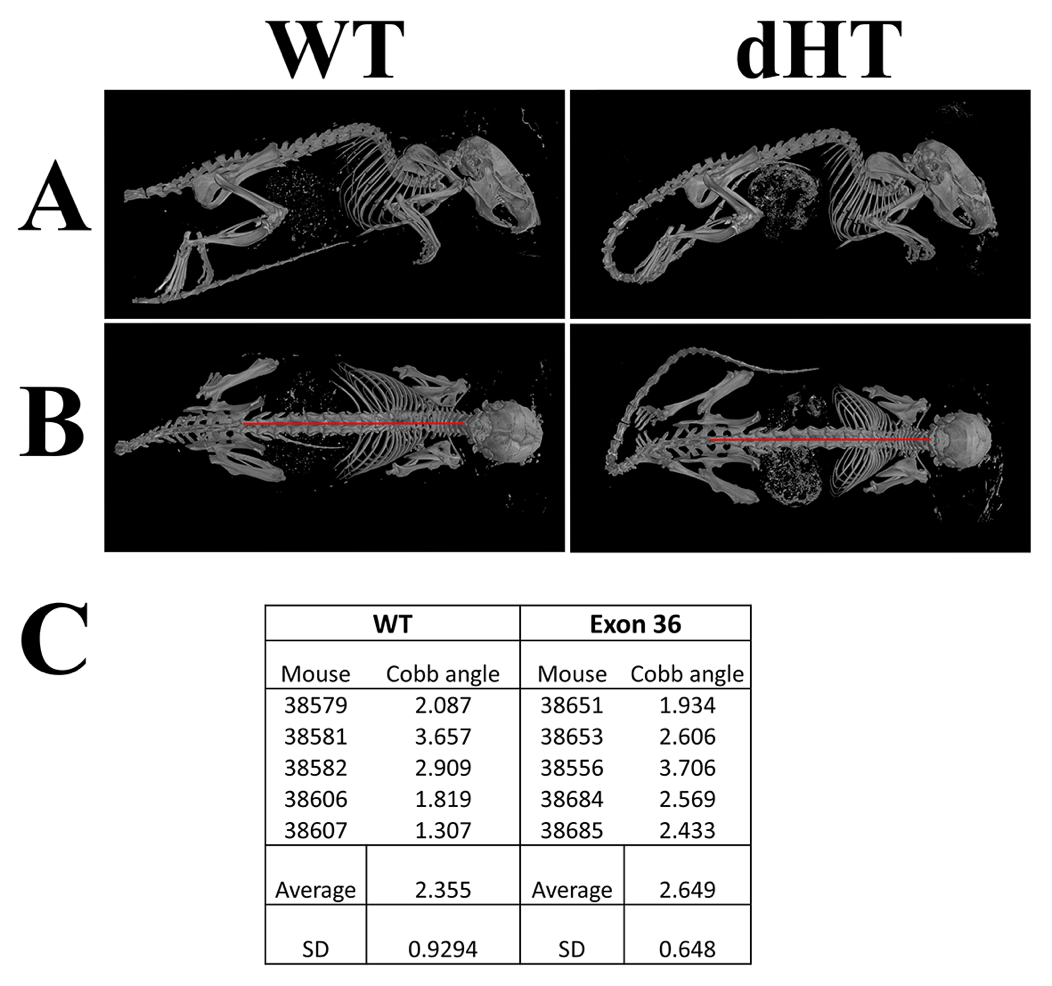

**Supplementary Figure 6: Ex36 mice do not show skeleton deformities nor scoliosis.** mCT imaging of skeletons from 12 weeks old WT and Ex36 mice **A.** Frontal view and **B**. top view of the whole-body skeleton of a WT (left) and Ex36 (right) littermate. The red line follows the spinal columns. **C.** Cobb angles were not different between the two mouse genotypes.

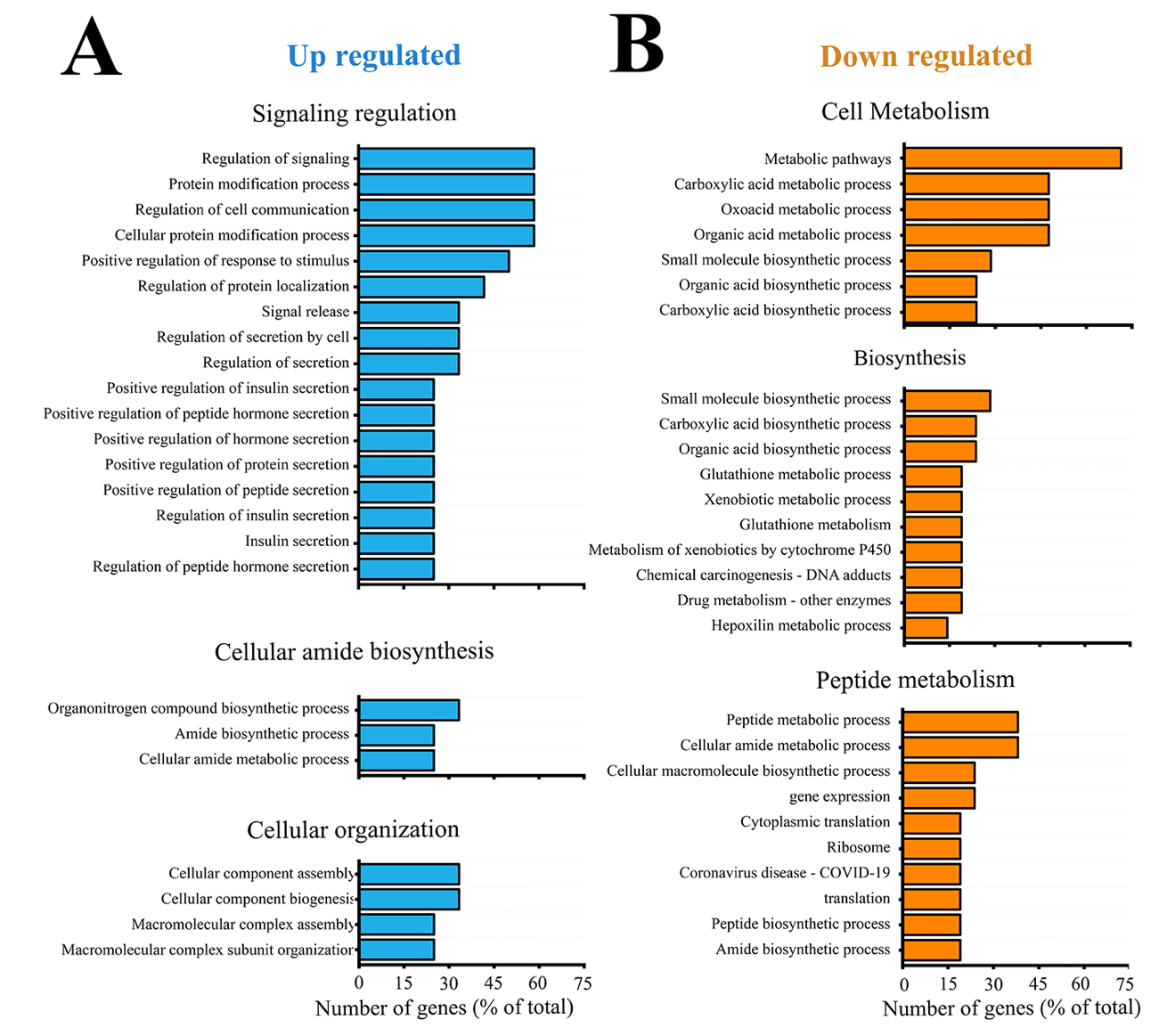

**Supplementary Figure 7**: **GO analysis of proteins showing a significant change in content in intrafusal fibers from dHT compared to WT littermates**. **A**. Proteins showing a significant enrichment in dHT versus WT intrafusal muscles were analyzed by GO Pathway analysis. The number of genes annotated to each cluster was calculated as a percentage over the total number of upregulated genes (=14). **B.** Proteins showing a significant depletion in dHT versus WT intrafusal muscles were analyzed by GO Pathway analysis. The number of genes annotated to each cluster was calculated as a percentage over the total number of down-regulated genes (=24).
